# Motion of single molecular tethers reveals dynamic subdomains at ER-mitochondria contact sites

**DOI:** 10.1101/2022.09.03.505525

**Authors:** Christopher J. Obara, Jonathon Nixon-Abell, Andrew S. Moore, Federica Riccio, David P. Hoffman, Gleb Shtengel, C. Shan Xu, Kathy Schaefer, H. Amalia Pasolli, Jean-Baptiste Masson, Harald F. Hess, Christopher P. Calderon, Craig Blackstone, Jennifer Lippincott-Schwartz

**Author notes:** These authors contributed equally to this work (Christopher J. Obara, Jonathon Nixon-Abell). Present Addresses: J.N-A., Cambridge Institute for Medical Research (CIMR), Cambridge, UK; F.R., Centre for Gene Therapy & Regenerative Medicine, King’s College London, London, UK; D.P.H., 10x Genomics, Pleasanton, CA, USA; C.S.X., Department of Cellular and Molecular Physiology, Yale University School of Medicine; C.B., MassGeneral Institute for Neurodegenerative Disease, Massachusetts General Hospital, Charlestown, MA, USA and Department of Neurology, Massachusetts General Hospital and Harvard Medical School, Boston, MA, USA. Correspondence (J.L-S.), (C.J.O). Correspondence and requests for materials should be addressed to Jennifer Lippincott-Schwartz, or Christopher Obara.

## Abstract

To coordinate cellular physiology, eukaryotic cells rely on the inter-organelle transfer of molecules at specialized organelle-organelle contact sites^1,2^. Endoplasmic reticulum-mitochondria contact sites (ERMCSs) are particularly vital communication hubs, playing key roles in the exchange of signaling molecules, lipids, and metabolites^3^. ERMCSs are maintained by interactions between complementary tethering molecules on the surface of each organelle^4,5^. However, due to the extreme sensitivity of these membrane interfaces to experimental perturbation^6,7^, a clear understanding of their nanoscale structure and regulation is still lacking. Here, we combine 3D electron microscopy with high-speed molecular tracking of a model organelle tether, VAPB, to map the structure and diffusion landscape of ERMCSs. From EM reconstructions, we identified subdomains within the contact site where ER membranes dramatically deform to match local mitochondrial curvature. In parallel live cell experiments, we observed that the VAPB tethers that mediate this interface were not immobile, but rather highly dynamic, entering and leaving the site in seconds. These subdomains enlarged during nutrient stress, indicating ERMCSs can readily remodel under different physiological conditions. An ALS-associated mutation in VAPB altered the normal fluidity of contact sites, likely perturbing effective communication across the contact site and preventing remodeling. These results establish high speed single molecule imaging as a new tool for mapping the structure of contact site interfaces and suggest that the diffusion landscape of VAPB is a crucial component of ERMCS homeostasis.

---

The most prevalent sites of contact between organelles in mammalian cells are between the endoplasmic reticulum (ER) and mitochondria^8,9^. This interface has been implicated in a multitude of biological processes in both health and disease, ranging from lipid synthesis and catabolism to calcium signaling and facilitation of cellular respiration^2,3,10^. Nevertheless, technical limitations have prevented the understanding of many aspects of ER-mitochondria contact site (ERMCS) biology (see Supplementary text for discussion)^6^, including the dynamic regulation of their molecular components and substructure. Here, we introduce quantitative imaging approaches that combine 3D electron microscopy and live cell, high-speed single molecule imaging to describe the substructure and dynamics of molecules within ERMCs under different physiological conditions.

To visualize the 3D architecture of ERMCs, we used high pressure freezing followed by freeze substitution and focused ion beam-scanning electron microscopy (FIB-SEM)^11^. This approach preserves contact sites in their near native state^7^, avoiding chemical fixation artifacts that might disrupt their delicate organization^12–14^. ER and mitochondria in small volumes from several COS7 cells were manually reconstructed, and regions of ER membrane within 24 nm of the outer mitochondria membrane were identified as sites of contact (Fig. 1a, Extended Data Fig. 1, Video 1). Highly abundant in the FIB-SEM volumes, ERMCSs were often near the base of mitochondrial cristae (Fig. 1b) or found at regions of constriction in the mitochondrial outer membrane (Extended Data Fig. 1), supportive of known roles of ERMCSs in mitochondria energy production and mitochondria fission^15–17^.

**Fig. 1.**
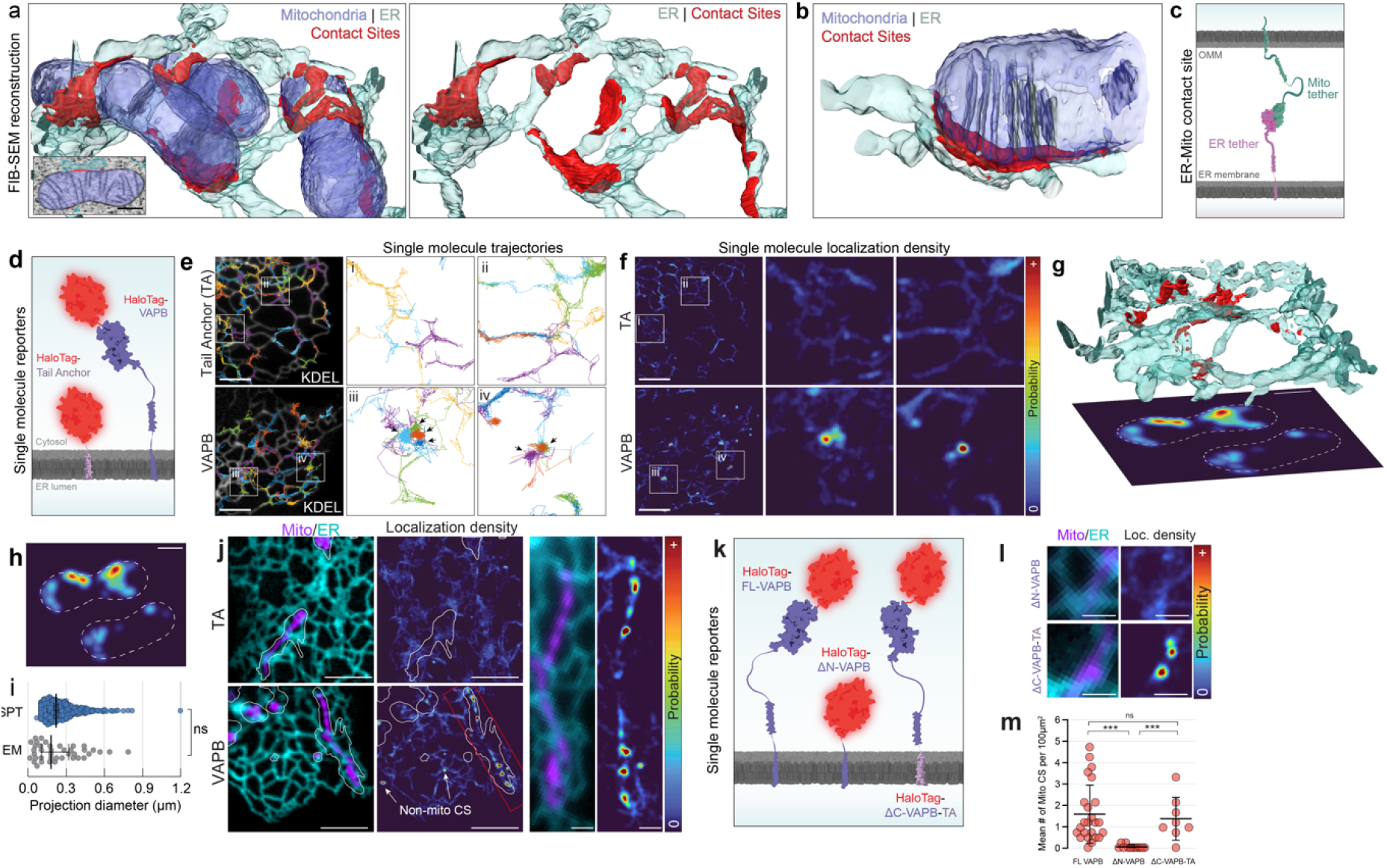
Contact sites between ER and mitochondria are visible as locations of altered ER tether motion. **a,** 3D EM reconstruction of ER (cyan) and mitochondria (blue) in a COS7 cell using data collected by FIB-SEM. ER membrane in the contact site is colored in red. Inset is a single slice showing voxel classifications. **b,** A single site of contact between the ER and a mitochondrion from (a) showing alignment of the mitochondrial cristae with the ER contact site. **c,** A cartoon schematic of a putative pair of ER-mitochondrial tethers. **d,** Cartoon schematic showing single molecule tracers used in e-f. **e,** sptPALM trajectories of single HaloTag molecules targeted to the ER membrane with a nonspecific tail anchor (TA) or as an N-terminal fusion on VAPB in COS7 cells. White is an ensemble luminal label for the ER (KDEL). Black arrows indicate regions of spatially correlated motion. **f,** The spatially-defined probability of finding a HaloTag label in the regions shown in e. Hot spots indicate likely sites of VAPB tethering. **g,** 3D EM reconstruction of the ER in association with two mitochondria (not shown). Contact sites are shown in red. Simulated sptPALM localization densities from the volume are shown below. Mitochondria positions outlined with dashed lines. **h,** Simulated localization densities from g. **i,** The size of contact sites as measured from FIB-SEM simulation projections or from single molecule localization density. **j,** Fluorescence micrograph of ER and mitochondria with the simultaneously acquired probability of TA or VAPB localization. White outlines indicate the space explored by the mitochondria within the 60 second time window used for the experiment. The inset is a zoom of the region in the red box. **k,** A cartoon schematic of the single molecule tracers used in l-m. **l,** Representative regions of contact between ER and mitochondria with associated localization densities are shown for the hemiprotein tracers shown in j. **m,** The number of mitochondria-associated regions of tracer tethering for each of the constructs in j-k. Scale bars: a,h,i,k; 500nm. e,f,j; 5 μm, insets, 1μm. l; 1μm.

With this view of ERMCS ultrastructure, we next sought to characterize ERMCSs at the molecular level in the dynamic setting of a living cell. Specific sites of contact between ER and mitochondria are mediated in *trans* by pairs of molecular tethers (Fig. 1c)^1,4,5^. Thus, we looked for locations where ER-localized tethers exhibited patterns of reduced single molecule motion, reasoning these may reveal the sites where they interact with their mitochondrial binding partners. We used single particle tracking-photoactivation localization microscopy (sptPALM)^18^ to follow the motion of individual molecules, while simultaneously capturing the location of the ER to inform subsequent tracking steps and minimize artifacts in forming trajectories (Video 2, Supplementary Text). Single molecules in this system were visualized by genetically fusing them to a HaloTag and labeling them with a photoactivatable version of JF646 ^19^, which was photoconverted at very low efficiency to ensure single proteins were tracked correctly with minimal linking artifacts (Fig. 1d, Video 2).

In order to interpret patterns of molecular motion, we first used a probe that does not associate with contact sites, a HaloTag-targeted to the ER with a minimal tail anchor (HaloTag-TA). We found that it explored the surface of the ER randomly, showing no detectable preferences for any specific ER regions (Fig. 1e, Tail Anchor). In contrast, when we then tracked a well-established ER-localized mitochondrial tether, VAMP-associated protein B (VAPB)^20–22^, we observed clusters of highly spatially associated trajectories (Fig. 1e, VAPB, see black arrows). By converting all trajectories within a specific time window to a spatially defined probability function (see Methods), locations of spatially associated trajectories appeared as ER regions with elevated probability of VAPB localization (Fig. 1f). These probability “hot spots” were present in the VAPB dataset but not with Halo-TA (Fig. 1f), and they were consistent with the size of ERMCSs extracted from FIB-SEM volumes (Fig. 1g-i).

Simultaneous imaging of ER and mitochondria in conjunction with sptPALM (see Methods) confirmed that a majority of VAPB hotspots were located on regions of ER close to mitochondria (Fig. 1j, Video 2), consistent with hotspots being areas of increased ER-mitochondria tethering. Of note, some probability hotspots also occurred in regions of ER with no mitochondria nearby (Fig. 1j, arrows, Extended Data Fig. 2,), likely the result of VAPB’s known role as a tether for additional organelles^21,22^. The formation of VAPB hot spots was entirely dependent on the ability of the protein to act as a tether by binding its interaction partner through its N-terminal cytosolic domain. Truncation of this N-terminal domain abrogated hotspot formation, and conversely, fusion of the N-terminal domain to the ER-targeted TA control was sufficient to form hotspots (Figs. 1k-m). We thus concluded that VAPB-associated hotspots at mitochondria represented tether interactions at *bona fide* ERMCSs.

In addition to providing a super-resolved image of ERMCSs in living cells, our sptPALM-based approach permitted analysis of how single VAPB molecules behaved before and after contact site interaction (Fig. 2a-b). Even within a short time window, many different VAPB molecules explored each contact site. VAPB molecules resided within a contact site only briefly, with most molecules leaving the site within a few seconds (Extended Data Fig. 2, Video 3). Using a nonparametric Bayesian approach^23^, trajectories of sufficient length (>500 steps, ~5.5 seconds) were broken into segments that were estimated to represent distinct kinetic states^24^ (Extended Data Fig. 3, Supplementary Text). This allowed classification of segments that were either freely diffusing in the ER (blue) or contact site-associated (red) (Fig. 2a-b, Extended Data Fig. 3). When analyzed over many cells and contact sites, we found that single VAPB molecules in ERMCSs showed significantly reduced diffusion (*D*_eff_) relative to VAPB molecules in surrounding ER tubules (Fig. 2c). At the same time, VAPB molecules within ERMCSs retained signatures of diffusive motion compared to immobilized bead controls (Fig. 2d; Extended Data Fig. 3, see Supplementary Text), indicating that VAPB molecules within contact sites still underwent dynamic motion.

**Fig. 2.**
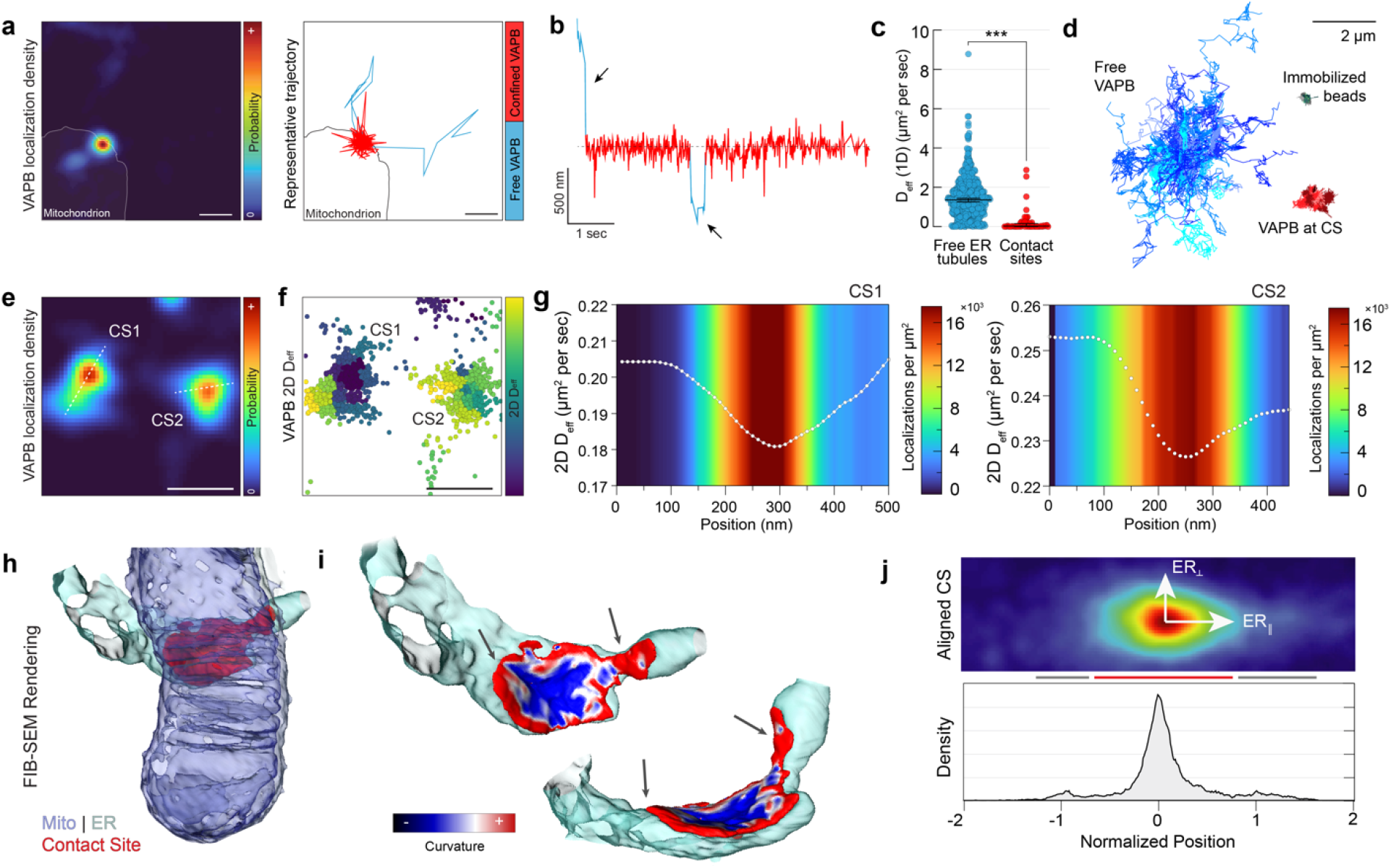
A variable VAPB diffusion landscape within single ER-mitochondria contact sites. **a,** The VAPB localization density of a single ER-mitochondria contact site and a representative trajectory of one VAPB molecule interacting with the site. Trajectory segments are color coded by the determined diffusion state. **b,** Time projection of the trajectory shown in a, color coded by diffusion state. Arrows indicate trajectory segments of free ER diffusion. **c,** Effective 1D diffusion coefficient of trajectory segments in ER tubules versus in mitochondria-associated contact sites. **d,** Grouped and overlayed representative trajectory segments of VAPB molecules associated with free ER diffusion, mitochondria-associated contact site diffusion, or immobilized beads. **e,** Localization density of two adjacent ER-mitochondria contact sites on the same mitochondrion. Lines represent axes chosen for plots in g. **f,** Location of single molecule steps associated with the contact sites in e, colored by the mean 2D D_eff_ in the local neighborhood. **g,** Mean 2D D_eff_ in a 30 nm neighborhood at each 10nm step along the lines in e. Color in background corresponds to the localization density. **h,** 3D EM reconstruction of a single ER-mitochondria contact site, ER (cyan), mitochondria (blue). ER membrane within the contact site is red. **i,** Local curvature of the ER membrane within the contact site. Arrows indicate regions of ER membrane with normal positive ER curvature that are within potential tethering distance for VAPB. **j,** Collective VAPB localization density across aligned mitochondria-associated contact sites on isolated ER tubules in the cellular periphery. The density shows an approximately Gaussian decay at the contact site center (red bar) and clear shoulders extending into the ER tubules (grey bars). Scale bars: a,b,e,f; 500 nm, time bar 1 sec. d; 2 μm.

Although some ERMCSs moved during the experiment or were short-lived (<1 min) (Extended Data Fig. 2), most remained stable over 60-90 seconds of imaging. During this time, many individual VAPB molecules repeatedly explored the same contact site interface. To map the diffusion landscape at this interface, we divided the contact site into small regions and analyzed the average diffusion of trajectory segments within each neighborhood (Extended Data Fig. 4, Supplementary Text)^25,26^. Examining the diffusive behavior of VAPB in each stable contact site in our dataset, we discovered a consistent pattern in VAPB diffusion behavior. Every contact site examined shared a common structure, with a central subdomain of low diffusion that gradually returned to the diffusion landscape of the surrounding ER at the edges of the contact site (Fig. 2e-g). The location and shape of this low diffusion subdomain closely correlated to the likelihood of finding a VAPB molecule (Fig. 2g), indicating that sites of slowed diffusion in the central subdomain were also elevated in tether abundance.

Since VAPB is believed to play an important role in modulating the structure of contact sites themselves^1,21^, we examined the ultrastructure of ER-mitochondria contact sites in our FIB-SEM dataset to identify any correlates to the low diffusion subdomains and associated hotspots of VAPB abundance seen with sptPALM. Focusing on the curvature of ER membranes at contact sites (see Supplementary Methods), we found that ER in each contact site had a distinct region of net negative curvature at its center (Fig. 2h-i). In this core region of the ERMCS, the ER adopted the inverse shape of the mitochondrion to which it was bound, suggesting adhesive forces at the site are sufficient to deform the ER membrane. Immediately flanking this negative curvature subdomain within the contact site were regions of ER membrane that retained normal positive curvature, despite being within tethering range of the mitochondrion (Fig 2i, arrows). Thus, adhesive forces in the contact site are likely not evenly spread throughout the structure, instead preferentially being enriched at the central subdomain.

Regions of negative ER curvature at the contact site center seen by FIB-SEM were reminiscent of the structure of subdomains of slowed VAPB diffusion and higher VAPB density observed by sptPALM. In order to directly compare these structures to one another, we aligned all sptPALM-detected contact sites localized along simple tubular ER structures to the direction of the incoming ER tubule. The summed probability density at these contact sites afforded increased resolution and showed that the distinct peak of VAPB intensity had clear probability shoulders extending into the surrounding ER tubule (Fig. 2j, gray bars, Extended Data Fig. 4), which closely resembled the pattern of negative/positive curvature areas within contact sites from FIB-SEM datasets (Fig. 2i). Thus, both structural FIB-SEM data and patterns of VAPB molecular diffusion support the presence of a central subdomain within ERMCSs that has higher VAPB abundance and greater adhesion properties. As VAPB probability density increased gradually toward this central ERMCS subdomain, contact sites may exist as metastable interfaces, in which many binding/unbinding events between tethers give rise to an overall adhesion force that holds the ER and mitochondria closely together.

To understand how the overall size of ERMCSs is regulated, we examined ERMCS size dependency on the availability of associated tethering molecules. The best characterized binding partner for VAPB in the mitochondria is the mitochondrial outer membrane component, protein tyrosine phosphatase interacting protein 51 (PTPIP51) ^21,27,28^ (Fig. 3a). We transiently overexpressed PTPIP51 in COS7 cells and performed sptPALM experiments. Contact site size significantly increased and covered large portions of the mitochondria compared to control cells not expressing PTPIP51 (Fig. 3b). The ERMCS expansion occurred across both axes of the structure (Fig. 3c-d). To test whether contact site expansion was equally dependent on both members of the tethering pair, we labeled the ER and mitochondria in COS7 cells and overexpressed either VAPB or PITPIP51, evaluating ER and mitochondrial morphology by Airyscan imaging. ER and mitochondrial morphology were insensitive to overexpression of VAPB at the time points assayed, but they showed dramatic rearrangements upon overexpression of PTPIP51, with the ER now completely enveloping most mitochondria (Fig. 3e). Despite this dramatic structural rearrangement, analysis of the sptPALM data showed no increase in the number of contact sites per cell. There was, however, a loss of nearly all VAPB contact sites associated with organelles other than mitochondria (Fig. 3f), suggesting that in this condition VAPB-binding at the mitochondria outcompeted its potential interactions at non-mitochondria associated sites. In agreement, cells highly overexpressing both VAPB and PTPIP51 showed essentially complete depletion of the ER pool of VAPB and selective enrichment to expanded contact sites (Fig. 3g). Thus, the availability of the mitochondrial side of the interaction controls VAPB-mediated tethering between ER and mitochondria.

**Fig. 3.**
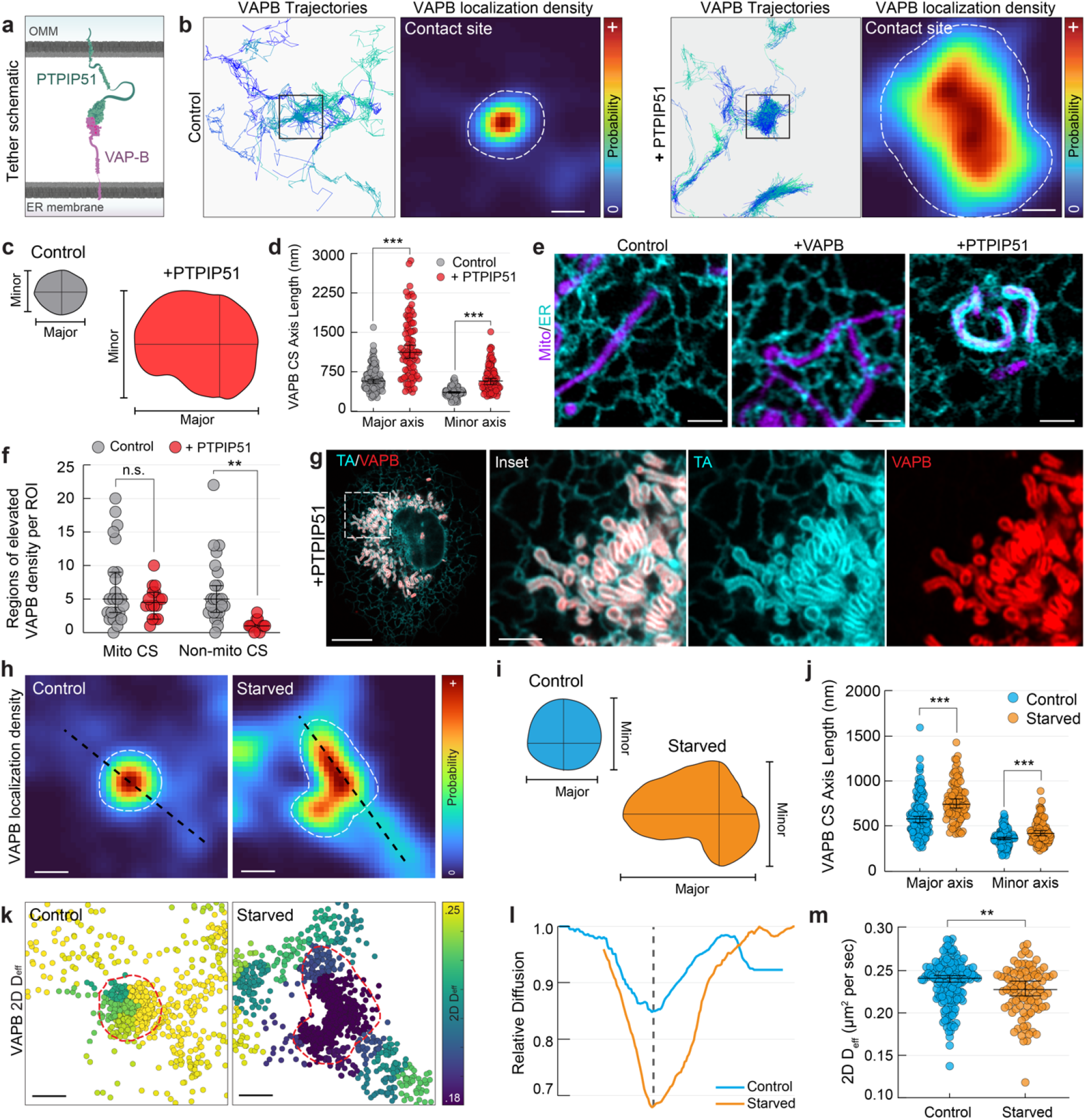
VAPB contact sites dynamically change size in response to tether availability and cellular metabolic needs. **a,** A cartoon showing the approximate size and orientation of VAPB and its OMM tether, PTPIP51. **b,** Representative trajectories of VAPB in the ER of COS7 cells or COS7 cells overexpressing PTPIP51. VAPB localization density is shown within the regions indicated, which are associated with mitochondria (not shown). Dotted lines delineate edge of the contact site. **c,** The contact sites in b with the approximate major and minor axes drawn as analyzed in d. **d,** The size of mitochondria-associated VAPB contact sites in the presence or absence of PTPIP51 overexpression. **e,** Representative Airyscan micrographs of the ER (cyan) and mitochondria (magenta) labeled with luminal markers in cells overexpressing VAPB or PTPIP51. **f,** The number of VAPB contact sites associated with mitochondria or other organelles in COS7 cells or COS7 cells overexpressing PTPIP51. **g,** An Airyscan micrograph showing localization of a fluorescent label targeted nonspecifically to the ER (TA) or VAPB in cells highly overexpressing PTPIP51. VAPB shows nearly complete depletion from the peripheral ER tubules. **h,** VAPB localization density of representative contact sites in COS7 cells cultured in complete medium (Control) or HBSS (Starved) for 8 hours. Lines are associated with the axes used for l. Dotted lines delineate edge of the contact site. **i,** Schematic of approximate major and minor axes for the contact sites in h. **j,** Contact site size in control or HBSS-starved cells. **k,** Single molecule steps associated with the contact sites in h, colored by the mean 2D D_eff_ in the local neighborhood. **l,** Effective 2D diffusion coefficient along the lines shown in h, normalized to the neighboring ER regions. The dashed line indicated the center of the contact site. **m,** Mean 2D D_eff_ within contact sites in COS7 cells under fed conditions or after 8 hours of HBSS-starvation. Scale bars: b,h,k; 200 nm. e; 2.5 μm. g; 10 μm, 2.5μm.

We explored how ERMCS structure and dynamics adapt to particular physiological challenges for meeting cellular needs. For example, previous biochemical and cryo-EM studies have suggested ERMCSs expand and change composition during acute nutrient deprivation^10,29–32^. These changes are proposed to provide crucial lipids and calcium to mitochondria to fuel oxidative phosphorylation or stimulate apoptosis^33,34^. To investigate this expansion and its effects on contact site substructure, we employed our sptPALM approach. VAPB trajectories were used to measure the size and organization of contact sites in COS7 cells after 8 hours of acute nutrient deprivation in HBSS. A significant expansion in the size of contact sites was observed (Fig. 3h-j, Extended Data Fig. 5). In addition, individual VAPB molecules still entered and left a contact site in seconds. This indicated that VAPB molecules in starved cells are not immobilized in contact sites; rather, the sites retain a fluid character. Mapping the diffusion landscape of VAPB in contact sites within starved cells, we observed that they still contained a central low diffusion subdomain despite their expanded size and shape (Fig. 3k-l, Extended Data Fig. 5). *D*_eff_ of VAPB within the central subdomain, however, was significantly lower than that measured in contact sites from well-fed cells (Fig. 3l-m). Thus, ERMCSs are capable of remodeling in response to changing metabolic needs. In the case of starvation, the expanded ERMCS interface, with its larger centralized hub of VAPB enrichment, likely facilitates more efficient metabolite exchange between ER and mitochondria to enable the cell to adapt to this stress condition.

Separately, we examined whether disease-causing forms of VAPB affected contact site organization. Several heterozygous mutations in the gene encoding VAPB cause the motor neuron disease known as amyotrophic lateral sclerosis (ALS) in humans^35^. In patients, ALS is most clearly associated with hyper-functional mitochondria and associated oxidative stress^36,37^. Disease-causing VAPB mutations generally encode more aggregation-prone versions of the protein and confer disease as a dominant, highly penetrant allele^38^. Aggregated VAPB molecules are nonfunctional^39–42^ and degrade quickly^43^, so disease states have been proposed to be conveyed through toxic effects of aggregates or by gene dosage effects due to aggregated proteins being degraded^38^. An additional possibility, however, is that prior to becoming immobilized in aggregates and eventually degraded, mutant VAPB molecules in the ER target to ER-mitochondrial contact sites and cause aberrant contact site function and/or dynamics.

To test the feasibility of this latter possibility, we examined whether VAPB molecules carrying ALS-causing mutations still target to ER-mitochondria contact sites and disrupt their organization. We tracked the motion of single VAPB molecules carrying the well-characterized ALS8 missense mutation, P56S, found in the MSP domain responsible for mitochondria interaction (Fig. 4a)^21,44^. In agreement with results from biochemical studies describing insoluble ER-associated aggregates^39,45,46^, many P56S VAPB molecules were in an immobilized form (Fig. 4b-c), rarely changing to other states. A significant fraction of P56S VAPB molecules also showed signatures of free ER diffusion like WT VAPB (Fig. 4b-c), indicating that some VAPB molecules are still effectively inserted into the ER in an unaggregated form. Notably, however, an additional pool of P56S VAPB molecules showed distinct signatures of contact site-associated motion (Fig. 4b-c). This raised the possibility that VAPB P56S might impair ERMCS structure and/or dynamics.

**Fig. 4.**
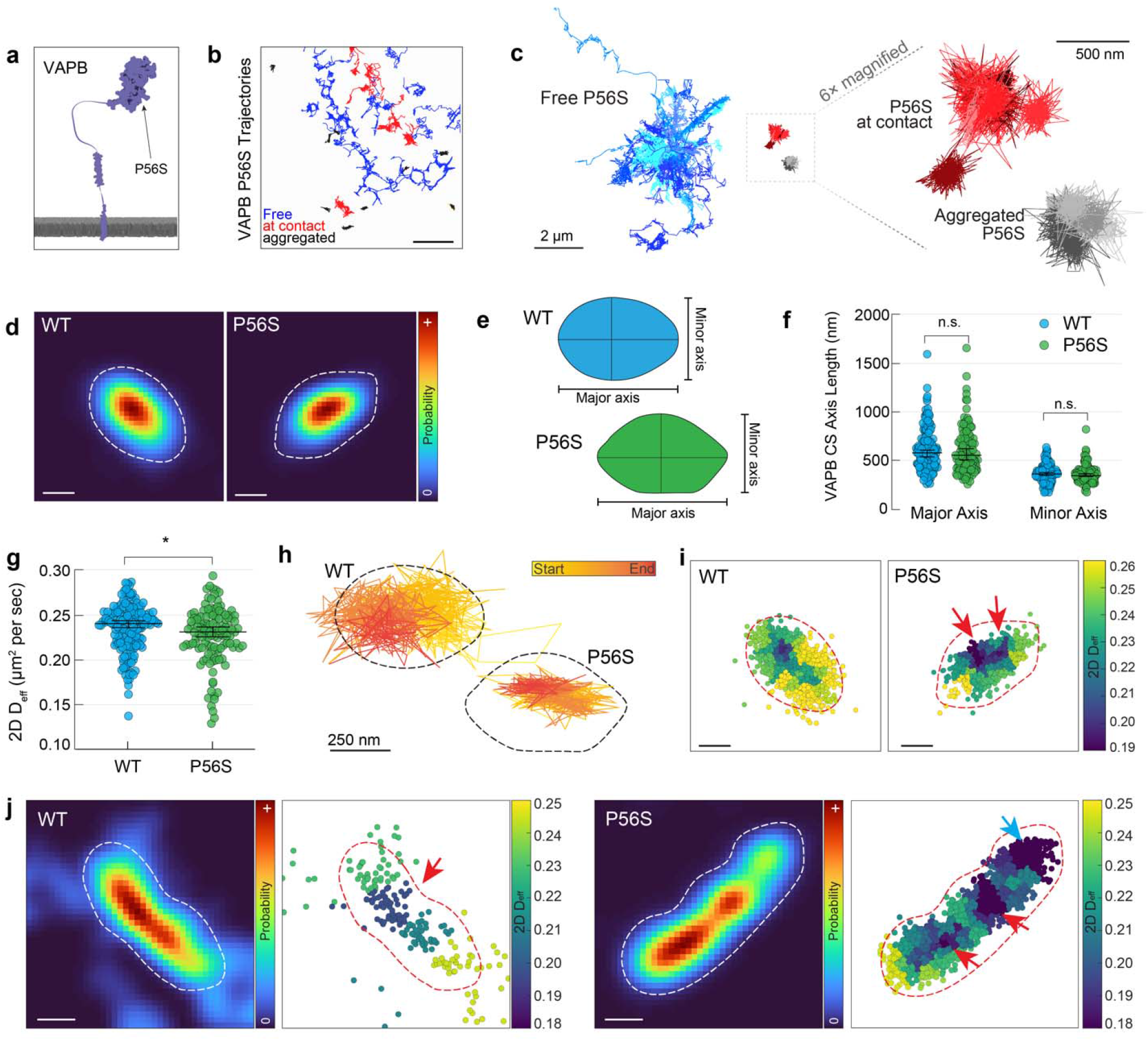
The ALS-linked VAPB P56S mutation displays aberrant motility at ER-mitochondria contact sites. **a,** A cartoon indicating the approximate location of the P56S mutation in the MSP domain of VAPB. **b,** Selected molecular trajectories of P56S VAPB in a representative COS7 cell. Tracks are color coded by their primary state of motion (see Methods). **c,** Grouped and overlaid representative trajectory segments of P56S VAPB in COS7 cells. **d,** Localization density in a representative ER-mitochondria contact site of HaloTag fused to either WT or P56S VAPB. Dotted lines delineate the edge of the contact site. **e,** Schematic illustrating dimensions of the contact sites in d. **f,** The size of contact sites as observed from trajectories of WT VAPB or P56S VAPB in COS7 cells. **g,** Mean 2D D_eff_ of WT VAPB or P56S VAPB within contact sites. **h,** A single, representative VAPB trajectory plotted from each of the contact sites shown in d. The boundary of the contact site is indicated with the dotted lines, color coding indicates the timing of the trajectory. **i,** Single molecule steps associated with the contact sites in d, colored by the mean 2D D_eff_ in the local neighborhood. Arrows indicate the asymmetric central diffusion well. **j,** Examples of localization densities of unusually elongated contact sites in the cellular periphery as seen with tracers of WT VAPB or P56S VAPB. Single molecule steps associated with the contact sites are displayed alongside. Arrows indicate the locations of distinct diffusional wells within the contact site. Scale Bars: b; 5 μm. c; 2 μm, inset 500 nm. d,h,j; 200 nm. i; 250 nm.

To assess whether ER-localized P56S VAPB molecules impact ERMCSs, we compared the properties of contact sites traced out by VAPB trajectories in cells expressing either WT or P56S VAPB. The overall localization densities of WT or P56S VAPB at contact sites were similar (Fig. 4d), and no significant differences were seen in contact site size or shape (Fig. 4e-f). However, the mean contact site associated *D*_eff_ was lower for P56S VAPB compared to WT VAPB (Fig. 4g), and there were significant differences in several features of their diffusion landscapes. Unlike WT VAPB, P56S VAPB molecules often did not explore the entire contact site area (Fig. 4h). Instead, they frequently showed a high degree of confinement in small subdomains, with P56S VAPB molecules trapped in these subdomains and unable to efficiently reach the edges of the contact site (Fig. 4h). The P56S VAPB confinement zones also correlated to regions of very low effective diffusion in the contact site landscape (Fig. 4i, see Supplementary Text). This was especially apparent for larger contact sites, in which multiple low diffusion subdomains of P56S VAPB could be seen dispersed across the structure, many of which did not correlate with areas of increased VAPB density (Fig. 4j, blue arrowhead). Thus, P56S VAPB molecules localized to ERMCSs generate aberrant subdomains with confined pools of P56S VAPB, and these subdomains are not the result of high tether density or the resulting negative curvature. Since trapped molecules do not exchange with the surrounding ER efficiently, this could lead to a more stable contact site interface, impacting contact site functionality.

## Discussion

Existing models of ERMCSs generally depict stable protein complexes tethering ER and mitochondria together. Our results combining high-speed molecular tracking with volumetric FIB-SEM paint a picture of ERMCSs that is much more complex. We discovered ERMCSs are exceptionally dynamic and highly organized, operating as a continuously changing interface that maintains spatially organized subdomains in the steady state. These metastable structures were adaptable, altering their size and configuration in response to specific physiological stimuli. This surprising plasticity likely is important in enabling ERMCSs to act as modular communication hubs for coordinating cell physiology.

Specifically, our high-speed molecular tracking revealed that ER-localized VAPB tethers diffused in and out of ERMCSs in seconds, exhibiting distinct patterns of slower motion when directly engaged with the contact site. Removal of VAPB’s mitochondrial interaction motif abolished the behavior, while overexpression of VAPB’s mitochondrial binding partner PTPIP51 exaggerated it. The affinity of VAPB for FFAT-containing binding partners like PTPIP51 is very low^21,47,48^, suggesting contact site interaction may consist of very many rapid binding and unbinding events across the structure. Consistent with this, we found a gradual increase in the VAPB abundance across the contact site with a clear peak at the center, where the likelihood of rebinding events would be elevated due to an increased density of potential tethering molecules within binding distance. Additionally, this higher abundance of tethers towards the center of the contact site would be predicted to provide increased adhesive force between the two membranes, a phenomenon we could directly observe as regions of negative curvature in the center of ERMCSs in FIB-SEM datasets. The surrounding edge regions of the contact site could serve as important staging regions for other ERMCS-associated activities. Supporting this possibility, the location of ERMCSs directly correlated to the location of mitochondrial cristae and sites of constriction in the mitochondrial membrane, biological phenomena that require recruitment of specific cellular machinery in addition to tethers^15–17,49,50^.

Our further results revealed ERMCS interfaces exhibited plasticity, able to adjust their structure and organization to different conditions. The size and shape of contact sites, for example, could be regulated through the availability of tethering partners like PTPIP51. Moreover, they became larger and exhibited reduced VAPB mobility during nutrient deprivation, which likely allows more efficient trans-organelle metabolite transfer, as has been previously reported in biochemical studies of cells undergoing acute starvation^29–32^. One way these contact site transformations might occur physiologically is through posttranslational modification of either ER- or mitochondria-localized ERMCS components by phosphatases and kinases^21,47,48,51,52^. Since VAPB rapidly exchanges between ER and ERMCS, VAPB’s posttranslational modifications could occur outside the contact site, where cytoplasmic enzymes would have easy access, a mechanism that could be generic for other tethers. The resulting adjustments in the size and shape of the contact site interface could create a unique membrane environment to stabilize other contact site molecules that are needed to accomplish the required metabolic adaptations.

The significance of contact site plasticity was further underscored by our high-speed tracking of P56S VAPB. Although most VAPB molecules with the P56S mutation were aggregated or undergoing free ER diffusion, a small but significant portion were still at ERMCSs. Notably, these ERMCS-associated P56S VAPB molecules disrupted normal VAPB diffusion in the contact site, with many molecules becoming trapped in multiple small subdomains of the ERMCS. We speculate that the inability of these trapped molecules to leave the ERMCS may lead to the impaired ability of the contact site to undergo normal dynamic restructuring. In agreement with this, more stable interfaces between ER and mitochondria have been shown in a variety of contexts to result in elevated mitochondrial function as a result of ER-to-mitochondria signaling^10,29,51,53^. Thus, even a small fraction of P56S in ERMCSs could directly lead to the oxidative stress known to occur in VAPB-associated ALS disease ^27,36,53^.

In summary, our results demonstrate that ERMCSs have distinct subdomains and a profoundly dynamic nature, with tethers diffusing into and out of the site over time scales of seconds and contact site size and configuration changing in response to different stimuli. Future work combining FIB-SEM structural data with single particle tracking promises to yield further insight into ERMCS biology, as well as impact understanding at the molecular level of other dynamic interfaces involved in inter-organelle communication.

Supplementary Information is available for this paper.

## Acknowledgements

We thank Carolyn Ott, Lorena Benedetti, Prabs Sengupta, Eric Betzig, and Pengli Zheng for advice on experimental protocols and critical reading of the manuscript. We thank Erik Snapp, Luke Lavis, the Janelia Open Chemistry and Tool Translation Teams for generously providing reagents and advice on their use. We are thankful to Eric Wait for computational support and the providing access to the HIP before public release.

## METHODS

### Coverslip cleaning and preparation

For single molecule imaging, high tolerance, Number 1.5 coverslips were purchased from Warner scientific (25mm) and precleaned with a modified version of a previously described protocol^1,2^. Coverslips were sonicated overnight (~12 hours) in 0.1% Hellmenex™ (Sigma), followed by five washes in 300 ml of distilled water. Coverslips were then transferred to a clean chamber of 300 ml of distilled water and sonicated overnight again, followed by five more washes. Coverslips were then ethanol sterilized in pure, 200 proof ethanol and air dried in a clean tissue culture hood. After cleaning, coverslips were stored in an airtight container until use, and were discarded if not used within 30 days of cleaning.

For FIB-SEM, sapphire coverslips (3mm diameter, 50μm thickness, Nanjing Co-Energy Optical Crystal Co. Ltd., COE) were cleaned for at least one hour in a basic piranha solution (5:1:1 water: ammonium hydroxide: hydrogen peroxide) followed by several washes in distilled water. The bottom of each coverslip had a thin layer of gold sputtered onto the side regions of the coverslip to distinguish top from bottom in subsequent steps (sputter coater Desk II, Denton Vacuum). After sputtering, coverslips were rinsed several times in distilled water and stored under vacuum in a desiccation chamber until use.

### Plasmids and reagents

ER-mRFP (Addgene #62236), mTagRFP-T2-Mito-7 (Addgene #58041) (referred to as mitoRFP in the text), mTagBFP2-N1 (Addgene #54566), mEGFP-N1 (Addgene #54767), mEGFP-C1 (Addgene #54759), and mEmerald-Sec61b-C1 (Addgene #90992) have been described previously, and were gifts from Erik Snapp, Michael Davidson, or generated in house. EGFP-VAPB^3^, HA-PTPIP51^4^, and pHAGE-Tet-STEMCCA^5^ have been previously described and were gifts from Pietro De Camilli, Kurt De Vos, and Robert Tijan, respectively.

All insertions and cassette changes were performed using the NEBuilder implementation of Gibson Assembly (New England Biolabs) unless specified otherwise, taking care to leave appropriate restriction sites for later changes. All constructs were sequenced before use and will be available on Addgene unless prohibited by copyright. Specific strategies and resulting plasmid maps are linked in the supplement.

### Cell culture and transfection

COS7 cells were purchased from ATCC and used within 40 passages. Cells were maintained in complete DMEM (phenol red-free Dulbecco’s modified Eagle medium (Corning) supplemented with 10% (v/v) fetal bovine serum (Corning), 2mM L-glutamine (Corning), 100U/ml penicillin and 100μg/ml streptomycin (ThermoFisher)). Cells were cultured at 37°C in 5% CO_2_, passaging was accomplished with 0.25% (w/v) trypsin (Corning) and care was taken never to let cells grow to more than 85% confluency or be seeded at less than 25% confluency, as they often become less flat after this.

For single molecule imaging, coverslips were precoated with 500μg/ml phenol red-free Matrigel depleted for growth factors (Corning) for one hour before plating in a 35mm tissue culture dish. Cells were seeded to ensure less than 60% confluency at the time of imaging in order to maximize regions of ER within the focal plane of the objective when focused just above the coverslip. Transfections were performed after letting the cells adhere to the glass for at least 12 hours using Fugene6 (Promega) at a 3:1 Fugene (ul) to DNA (ug) ratio according to the manufacturer’s protocol. Each 35mm dish was transfected with 1.5 ug of DNA using the following ratio: 750 ng PrSS-mEmerald-KDEL (ER structure label), 500 ng mito-dsRed (mitochondria structure label), and 250 ng of the HaloTag construct used for sptPALM, except for cells transfected with PTPIP51 which were given an extra 500ng of PTPIP51-IRES-mTagBFP2 in addition. Imaging was always performed at least 12 hours after transfection, but always before 24 hours and taking care to avoid cells showing morphological changes from ER and mitochondria label expression that become increasingly common at later time points. For starvation experiments, cells were incubated in HBSS for the last 8 hours before imaging, but complete medium was replaced on the cells immediately before imaging.

Cells prepared for Airyscan imaging were plated in high tolerance commercially acquired 35mm imaging chambers (MatTek Life Sciences). Briefly, coverslips were precoated with 500μg/ml phenol red-free Matrigel depleted for growth factors (Corning) for 20 min. Cells were then transfected in solution using Fugene6 (Promega) according to a modified version of the manufacturer’s protocol. Briefly, cells were resuspended after pelleting in prepared transfection complexes made according to the manufacturer’s recommendations in OptiMEM (ThermoFisher). Cells were incubated for 15 min at 37°C before being plated on the coated coverslips in 2 ml of complete Medium. Imaging was performed 18-24 hours post transfection.

### Halo Labeling

Cells were labelled for sptPALM by incubating with 10 nM PA-JF646^6^ in OptiMEM (ThermoFisher) for 1 minute followed at least 5 washes with 10 ml of PBS, performed while simultaneously aspirating and taking care to never let the cells come in direct contact with the air. The cells were then washed once with 10 ml of complete DMEM and left to recover in 2 ml of complete DMEM for 15 minutes before imaging.

Cells were labeled for Airyscan imaging by replacing the complete medium on the cells with complete DMEM supplemented with 10nM JF635^7^. The highly fluorogenic nature of this JF dye compound removes the need for washing steps, and the sample can directly be imaged on the microscope.

### Microscopy and imaging conditions

Single molecule imaging was performed using a custom widefield microscope assembled in an inverted Nikon Ti-E outfitted with a stage top incubator to stabilize temperature, CO_2_, and relative humidity during imaging (Tokai Hit). The flat lamella of cells where sptPALM is possible (approximately 500nm thick or less) were located using eye pieces to visualize the ER and mitochondria localization. To avoid bias, the experimenter was always blinded to the single molecule tracers. In experiments where PTPIP51-IRES-mTagBFP2 was overexpressed, the cells were selected using the fluorescence of the mTagBFP2 in addition to the ER and mitochondria structure.

Excitation was performed using three fiber-coupled solid state laser lines (488nm, 561nm, 642nm; Agilent Technologies) introduced into the system with a conventional rear-mount TIRF illuminator. The angle of incidence was manually adjusted for each cell beneath the critical angle to maximize the evenness of the illumination in the ER. The illumination in the 488nm and 561nm channel was manually adjusted based on the brightness of the sample to minimize fluorescent bleed through, but the total power on each line was always kept less than 50μW and 150μW total in the back aperture, respectively. Single molecules were always imaged using a constant total power of 11.5mW of 647nm light in the back aperture. If necessary, a small amount of 405 nm light was introduced to tune the photoactivation rate of the molecules being tracked, but in practice this was rarely needed.

Emitted light was collected with a 100x *α*-plan apochromat 1.49 NA oil immersion objective (Nikon Instruments) and focused through a MultiCam optical splitter (Cairn Research). The emission path was split onto three arms of the splitter using a 565LP and a 647LP dichroic mirror (Chroma) placed sequentially in the optical path to split the light from the 488nm and 561nm channels, respectively. These emission paths were additionally cleaned up by passing the emitted light through a 525/50 BP and a 605/70 BP filter (Chroma), respectively. The remaining light transmitted through the MultiCam represents the far-red signal where the single molecules of HaloTag-linked dye are imaged, and the path was passed through an additional 647LP filter to clean up any stray light in the system that could decrease the resolving power of the sptPALM approach. All three channels were collected from electronically synchronized iXon3 electron multiplying charged coupled device cameras (EM-CCD, DU-897; Andor Technology). In order to image quickly enough, the field of view was reduced to a 128 x 128 pixel square (20.48μm x 20.48μm). The location of the square was carefully chosen for each sample to contain the flattest region of ER possible while remaining near the center of the camera chip, since the objective in use is chromatically corrected to high precision only near the center of the field of view. Imaging was performed with 5 ms exposure times for 60-90 seconds at a time, and the timing of each frame was monitored using an oscilloscope directly coupled into the system (mean frame rate ~ 95Hz).

Airyscan imaging was performed using a commercially acquired Zeiss LSM 880 microscope with a live cell incubation system (Zeiss Microscopy). Briefly, labeled samples were sequentially excited with laser lines at 633nm, 561nm, and 488nm. Emission fluorescence is collected using a 63x 1.4 NA oil immersion objective (Zeiss Microscopy) with an open pinhole and passed through an appropriate custom bandpass filter based on the expected emission profile of the sample to the arrayed detector for the Airyscan unit (561nm,488nm: BP495-550 + LP570, 633nm: BP570-620 + LP645). Airyscan reconstruction and deconvolution was performed using the default settings (filter size=6). Images were pseudocolored and prepared using Fiji (NIH) for visibility.

### Channel registration and spectral analysis

At the time of imaging, a crude channel alignment was performed using a sparse distribution of Tetraspek beads on a coverslip, prepared and imaged as for other sptPALM samples. The angle of the dichroic mirrors was manually adjusted to get as much overlap between the channels in the main field of view as possible. In practice, this alignment was sufficient to support the manual steps in the tracking pipeline (see below), but applications requiring more precise alignment were accomplished using a custom subpixel alignment pipeline in Fiji.

Variation in expression level of the three markers (ER structure marker, mitochondria structure marker, single molecule tracer) due to uneven transfection is not a trivial issue, and often required manual adjustment in the relative laser power for the 488nm and 561nm lines by the experimenter. Since this could in principle create artifacts in the automated analysis pipeline or introduce erroneous single molecule localizations as a result of bleed through, all of the samples were run through an automated spectral analysis pipeline that checked for fluorescence contamination from the blue-shifted channels. Any samples where the detectable bleed-through contamination was significant compared to the signal of single molecules were removed before downstream analysis was performed.

### Localization and tracking

Localizations were identified in the single molecule datasets using a previously described pipeline^8^ to estimate positions and precision of localization using an MLE-based fitting approach. The quality filter used in the downstream tracking pipeline limited analyzable localizations to those identified with precision (as estimated from the Cramér-Rao lower bound) in the range of 20-30 nm.

Trajectories were assembled from single molecule images using the TrackMate plugin in Fiji ^9,10^. Linking parameters were experimentally selected for each data set to minimize visible linkage artifacts as determined by eye. Resulting putative trajectories were then projected on to the simultaneously collected ER network structure, and manually curated to remove any trajectory linkages that are close in 2D but far from one another in the underlying organelle structure. This step proved crucial to assembling trajectories that moved within the structure without linkage artifacts. Resulting trajectories were exported from TrackMate and imported into MATLAB for subsequent analysis.

### Spatial density analysis and contact site identification

Spatial probability density is mapped by choosing the spatiotemporal boundaries of the data to be analyzed (x, y, t) and binning the resulting localizations into 30nm square pixels. The resulting counts are normalized to the total number of localizations within the dataset, and as such probability represents solely the likelihood of a single molecule falling in a certain pixel if chosen at random (i.e., a spatially-defined probability mass function). This effectively minimizes the effects of differences in photoactivation efficiency or tagged protein expression level when identifying the boundaries of a contact site.

The initial location of contact sites was identified from the spatially defined probability density when calculated over the entire image, but the location and boundaries often had to be manually refined, especially under conditions where contact sites move or change orientation (Extended Data Fig. 3).

### Spatial clustering and diffusion landscape estimation

To generate a map of the diffusion landscape within contact sites, the space inside the contact site was divided into distinct compartments by Voronoi tessellation informed by the probability density at the site. The trajectories associated with the site were broken into single steps and assigned to a tessellation by the location of the beginning of the step^11,12^. Bayesian inference was then used to model the resulting distribution as an overdamped Langevin system within each tessellation, assuming single molecules in the same space at distinct times can be viewed as independent experiments probing the same molecular environment. The resulting diffusive component was reported for each tessellation as an effective 2D diffusion coefficient.

### Identification of latent states in single trajectories

All trajectories longer than 500 steps (~5.5 seconds) were analyzed using a nonparametric Bayesian modeling technique (Hierarchical Dirichlet Process Modeling, HDP) to estimate latent state changes in single molecule behavior^13,14^. Briefly, the system was treated as a switching linear dynamical system (SLDS). As in previous work^13,14^, an overdamped Langevin equation was used to interpret the parameters of the linear dynamical system used in the SLDS model. This approach removes the need for an upper bound on the number of potential states common in SPT analysis approaches (e.g., Hidden Markov Models, etc.), which become intractable in a spatially complex environment like the ER (see Supplementary Text). Eigen-decompositions of the implied force and diffusion tensors for each determined state enables 1D analysis through tensor diagonalization. Note this SLDS treats thermal fluctuations as a distinct component from measurement noise, allowing diffusive properties of the system to be quantitatively estimated independently of measurement noise (e.g. localization errors).

### High pressure freezing and freeze substitution

Immediately prior to freezing, cells were manually inspected using an inverted widefield microscope to ensure reasonable morphology and viability. Cells were then transferred to a water jacketed incubator (ThermoFisher, Midi 40) where they were kept at 37°C in 5% CO_2_ and 100% humidity until ready for freezing. Each sapphire coverslip was removed one at time from the incubator, overlaid with a 25% (w/v) solution of 40,000 MW dextran (Sigma), loaded between two hexadecane-coated freeing platelets (Technotrade International), and placed in the HPF holder. Freezing was then performed using a Wohlwend Compact 2 high pressure freezer, according to the manufacturer’s protocol. Frozen samples were stored under liquid nitrogen until freeze substitution was performed.

Freeze-substitution was performed with a modified version of a previously described protocol^15,16^. Briefly, frozen samples were transferred to cryotubes containing freeze-substitution media (2% OsO_4_, 0.1% Uranyl acetate, and 3% water in acetone) and placed in an automated freeze substitution machine (AFS2, Leica Microsystems). A freeze substitution protocol was used as previously described^16^, and the samples were then washed three times in anhydrous acetone and embedded in Eponate 12 (Ted Pella, Inc.). The sapphire coverslip was then removed and the block was re-embedded in Durcapan (Sigma) resin for FIB-SEM imaging.

### FIB-SEM

FIB-SEM was performed essentially as previously described^16–18^. Briefly, a customized FIB-SEM using a Zeiss Capella FIB column fitted at 90 degrees on a Zeiss Merlin SEM was used to sequentially image and mill 8nm layers from the Durcapan-embedded block. Milling steps were performed using a 15 nA gallium ion beam at 30 kV to generate two sequential 4nm steps. Data was acquired at 500kHz/pixel using a 2nA electron beam at 1.0 kV landing energy with 8nm xy resolution to generate isotropic voxels. Datasets were registered post-acquisition using a SIFT-based algorithm^19^.

### Voxel classification and surface determination

Although several automated segmentation protocols exist for reconstruction of organelles from FIB-SEM data^18,20^, we found that minor errors in voxel classification within the contact sites themselves obscured our ability to analyze the local curvature in sufficient resolution for our needs (see Supplementary Text for discussion). Consequently, we selected a few small volumes containing mitochondria and manually classified the voxels for the ER using Amira (ThermoFisher). We used a modified watershed algorithm to classify the mitochondrial membranes but performed a manual curation to remove artifacts. Potential contact sites on the ER surface were identified as regions of ER membrane within 24nm (approximately 3 pixels) of the mitochondrial outer membrane, as measured by dilation of the mitochondrial outer membrane.

### Triangulation, smoothing, and curvature analysis

Triangulated surfaces were fit to the voxel classifications using a marching cubes-based algorithm implemented directly in Amira. To avoid voxel-step artifacts in the surface, gaussian smoothing was applied to the voxel data using a local likelihood measure selected over a kernel size relevant for the expected curvature of the underlying membrane (see Supplementary Text). The resulting triangulated surfaces were rendered for use in the figures using Amira, and they serve as the scaffold for subsequent curvature analysis.

Mean local curvature of the ER was computed as a scalar field over the triangulated surface using a 20-layer neighborhood to fit a quadratic form along the two principal curvature axes. Note this is different from gaussian curvature, and the resulting value is negative in strictly concave regions and positive in regions that are strictly convex. Scalar fields were calculated and mapped using the curvature field module in Amira. Details are given in the Supplement.

## References

1. Prinz, W. A., Toulmay, A. & Balla, T. The functional universe of membrane contact sites. Nat Rev Mol Cell Bio 21, 7–24 (2019).

2. Phillips, M. J. & Voeltz, G. K. Structure and function of ER membrane contact sites with other organelles. Nat Rev Mol Cell Bio 17, 69–82 (2016).

3. Zung, N. & Schuldiner, M. New horizons in mitochondrial contact site research. Biol Chem 401, 793–809 (2020).

4. Eisenberg-Bord, M., Shai, N., Schuldiner, M. & Bohnert, M. A Tether Is a Tether Is a Tether: Tethering at Membrane Contact Sites. Dev Cell 39, 395–409 (2016).

5. Scorrano, L. et al. Coming together to define membrane contact sites. Nat Commun 10, 1287 (2019).

6. Giamogante, F., Barazzuol, L., Brini, M. & Calì, T. ER–Mitochondria Contact Sites Reporters: Strengths and Weaknesses of the Available Approaches. Int J Mol Sci 21, 8157 (2020).

7. Pezzati, R., Bossi, M., Podini, P., Meldolesi, J. & Grohovaz, F. High-resolution calcium mapping of the endoplasmic reticulum-Golgi-exocytic membrane system. Electron energy loss imaging analysis of quick frozen-freeze dried PC12 cells. Mol Biol Cell 8, 1501–1512 (1997).

8. Valm, A. M. et al. Applying systems-level spectral imaging and analysis to reveal the organelle interactome. Nature 546, 162–167 (2016).

9. Heinrich, L. et al. Whole-cell organelle segmentation in volume electron microscopy. Nature 599, 141–146 (2021).

10. Csordás, G. et al. Structural and functional features and significance of the physical linkage between ER and mitochondria. J Cell Biology 174, 915–921 (2006).

11. Xu, C. S. et al. An open-access volume electron microscopy atlas of whole cells and tissues. Nature 599, 147–151 (2021).

12. Sosinsky, G. E. et al. The combination of chemical fixation procedures with high pressure freezing and freeze substitution preserves highly labile tissue ultrastructure for electron tomography applications. J Struct Biol 161, 359–371 (2008).

13. Hoffman, D. P. et al. Correlative three-dimensional super-resolution and block-face electron microscopy of whole vitreously frozen cells. Science 367, (2020).

14. Hicks, M. L., Brilliant, J. D. & Foreman, D. W. Electron Microscope Comparison of Freeze-Substitution and Conventional Chemical Fixation of Undecalificied Human Dentin. J Dent Res 55, 400–410 (1976).

15. Huynen, M. A., Mühlmeister, M., Gotthardt, K., Guerrero-Castillo, S. & Brandt, U. Evolution and structural organization of the mitochondrial contact site (MICOS) complex and the mitochondrial intermembrane space bridging (MIB) complex. Biochimica Et Biophysica Acta Bba - Mol Cell Res 1863, 91–101 (2016).

16. Kozjak-Pavlovic, V. The MICOS complex of human mitochondria. Cell Tissue Res 367, 83–93 (2017).

17. Friedman, J. R. et al. ER Tubules Mark Sites of Mitochondrial Division. Science 334, 358–362 (2011).

18. Manley, S. et al. High-density mapping of single-molecule trajectories with photoactivated localization microscopy. Nat Methods 5, 155–7 (2008).

19. Grimm, J. B. et al. Bright photoactivatable fluorophores for single-molecule imaging. Nat Methods 13, 985–988 (2016).

20. Nishimura, Y., Hayashi, M., Inada, H. & Tanaka, T. Molecular Cloning and Characterization of Mammalian Homologues of Vesicle-Associated Membrane Protein-Associated (VAMP-Associated) Proteins. Biochem Bioph Res Co 254, 21–26 (1999).

21. Murphy, S. E. & Levine, T. P. VAP, a Versatile Access Point for the Endoplasmic Reticulum: Review and analysis of FFAT-like motifs in the VAPome. Biochim Biophys Acta 1861, 952–61 (2016).

22. Kors, S., Costello, J. L. & Schrader, M. VAP Proteins – From Organelle Tethers to Pathogenic Host Interactors and Their Role in Neuronal Disease. Frontiers Cell Dev Biology 10, 895856 (2022).

23. Calderon, C. Data-Driven Techniques for Detecting Dynamical State Changes in Noisily Measured 3D Single-Molecule Trajectories. Molecules 19, 18381–18398 (2014).

24. Calderon, C. P. & Bloom, K. Inferring Latent States and Refining Force Estimates via Hierarchical Dirichlet Process Modeling in Single Particle Tracking Experiments. Plos One 10, e0137633 (2015).

25. Beheiry, M. E. et al. A Primer on the Bayesian Approach to High-Density Single-Molecule Trajectories Analysis. Biophys J 110, 1209–15 (2016).

26. Masson, J.-B. et al. Mapping the energy and diffusion landscapes of membrane proteins at the cell surface using high-density single-molecule imaging and Bayesian inference: application to the multiscale dynamics of glycine receptors in the neuronal membrane. Biophys J 106, 74–83 (2014).

27. Stoica, R. et al. ER–mitochondria associations are regulated by the VAPB–PTPIP51 interaction and are disrupted by ALS/FTD-associated TDP-43. Nat Commun 5, 3996 (2014).

28. Vos, K. J. D. et al. VAPB interacts with the mitochondrial protein PTPIP51 to regulate calcium homeostasis. Hum Mol Genet 21, 1299–1311 (2012).

29. Bravo, R. et al. Increased ER–mitochondrial coupling promotes mitochondrial respiration and bioenergetics during early phases of ER stress. J Cell Sci 124, 2143–2152 (2011).

30. Sood, A. et al. A Mitofusin-2–dependent inactivating cleavage of Opa1 links changes in mitochondria cristae and ER contacts in the postprandial liver. Proc National Acad Sci 111, 16017–16022 (2014).

31. Theurey, P. et al. Mitochondria-associated endoplasmic reticulum membranes allow adaptation of mitochondrial metabolism to glucose availability in the liver. J Mol Cell Biol 8, 129–143 (2016).

32. Wu, W. et al. FUNDC1 regulates mitochondrial dynamics at the ER–mitochondrial contact site under hypoxic conditions. Embo J 35, 1368–1384 (2016).

33. Eisner, V., Picard, M. & Hajnóczky, G. Mitochondrial dynamics in adaptive and maladaptive cellular stress responses. Nat Cell Biol 20, 755–765 (2018).

34. Simmen, T. & Herrera-Cruz, M. S. Plastic mitochondria-endoplasmic reticulum (ER) contacts use chaperones and tethers to mould their structure and signaling. Curr Opin Cell Biol 53, 61–69 (2018).

35. Borgese, N., Iacomino, N., Colombo, S. F. & Navone, F. The Link between VAPB Loss of Function and Amyotrophic Lateral Sclerosis. Cells 10, 1865 (2021).

36. Barnham, K. J., Masters, C. L. & Bush, A. I. Neurodegenerative diseases and oxidative stress. Nat Rev Drug Discov 3, 205–214 (2004).

37. Mejzini, R. et al. ALS Genetics, Mechanisms, and Therapeutics: Where Are We Now? Front Neurosci-switz 13, 1310 (2019).

38. Borgese, N., Navone, F., Nukina, N. & Yamanaka, T. Mutant VAPB: Culprit or Innocent Bystander of Amyotrophic Lateral Sclerosis? Contact 4, 25152564211022516 (2021).

39. Teuling, E. et al. Motor Neuron Disease-Associated Mutant Vesicle-Associated Membrane Protein-Associated Protein (VAP) B Recruits Wild-Type VAPs into Endoplasmic Reticulum-Derived Tubular Aggregates. J Neurosci 27, 9801–9815 (2007).

40. Suzuki, H. et al. ALS-linked P56S-VAPB, an aggregated loss-of-function mutant of VAPB, predisposes motor neurons to ER stress-related death by inducing aggregation of co-expressed wild-type VAPB: Detailed characterization of VAPB/ALS8. J Neurochem 108, n/a–n/a (2010).

41. Kim, S., Leal, S. S., Halevy, D. B., Gomes, C. M. & Lev, S. Structural Requirements for VAP-B Oligomerization and Their Implication in Amyotrophic Lateral Sclerosis-associated VAP-B(P56S) Neurotoxicity*. J Biol Chem 285, 13839–13849 (2010).

42. Yamanaka, T., Nishiyama, R., Shimogori, T. & Nukina, N. Proteomics-Based Approach Identifies Altered ER Domain Properties by ALS-Linked VAPB Mutation. Sci Rep-uk 10, 7610 (2020).

43. Papiani, G. et al. Restructured endoplasmic reticulum generated by mutant amyotrophic lateral sclerosis-linked VAPB is cleared by the proteasome. J Cell Sci 125, 3601–3611 (2012).

44. Dudás, E. F., Huynen, M. A., Lesk, A. M. & Pastore, A. Invisible leashes: The tethering VAPs from infectious diseases to neurodegeneration. J Biological Chem 296, 100421 (2021).

45. Nishimura, A. L. et al. A Mutation in the Vesicle-Trafficking Protein VAPB Causes Late-Onset Spinal Muscular Atrophy and Amyotrophic Lateral Sclerosis. Am J Hum Genetics 75, 822–831 (2004).

46. Fasana, E. et al. A VAPB mutant linked to amyotrophic lateral sclerosis generates a novel form of organized smooth endoplasmic reticulum. Faseb J 24, 1419–1430 (2010).

47. Yeo, H. K. et al. Phospholipid transfer function of PTPIP51 at mitochondria-associated ER membranes. Embo Rep 22, e51323 (2021).

48. Mattia, T. D. et al. FFAT motif phosphorylation controls formation and lipid transfer function of inter-organelle contacts. Embo J 39, e104369 (2020).

49. Anand, R., Reichert, A. S. & Kondadi, A. K. Emerging Roles of the MICOS Complex in Cristae Dynamics and Biogenesis. Biology 10, 600 (2021).

50. Lee, H. & Yoon, Y. Mitochondrial Fission: Regulation and ER Connection. Mol Cells 37, 89–94 (2014).

51. Stoica, R. et al. ALS/FTD-associated FUS activates GSK-3β to disrupt the VAPB–PTPIP51 interaction and ER–mitochondria associations. Embo Rep 17, 1326–1342 (2016).

52. Yang, K. et al. The Key Roles of GSK-3β in Regulating Mitochondrial Activity. Cell Physiol Biochem 44, 1445–1459 (2017).

53. Hirabayashi, Y. et al. ER-mitochondria tethering by PDZD8 regulates Ca2+ dynamics in mammalian neurons. Science 358, 623–630 (2017).

## Methods References

1. Sengupta, P. et al. Probing protein heterogeneity in the plasma membrane using PALM and pair correlation analysis. Nat Methods 8, 969–975 (2011).

2. Sengupta, P., Jovanovic-Talisman, T. & Lippincott-Schwartz, J. Quantifying spatial organization in point-localization superresolution images using pair correlation analysis. Nat Protoc 8, 345–354 (2013).

3. Dong, R. et al. Endosome-ER Contacts Control Actin Nucleation and Retromer Function through VAP-Dependent Regulation of PI4P. Cell 166, 408–423 (2016).

4. Vos, K. J. D. et al. VAPB interacts with the mitochondrial protein PTPIP51 to regulate calcium homeostasis. Hum Mol Genet 21, 1299–1311 (2012).

5. Sommer, C. A. et al. Induced Pluripotent Stem Cell Generation Using a Single Lentiviral Stem Cell Cassette. Stem Cells 27, 543–549 (2009).

6. Grimm, J. B. et al. Bright photoactivatable fluorophores for single-molecule imaging. Nat Methods 13, 985–988 (2016).

7. Grimm, J. B. et al. A general method to fine-tune fluorophores for live-cell and in vivo imaging. Nat Methods 14, 987–994 (2017).

8. Li, Y. et al. Optimal 3D single-molecule localization in real time using experimental point spread functions. Nat Methods 15, 367–369 (2018).

9. Jaqaman, K. et al. Robust single particle tracking in live cell time-lapse sequences. Nat Methods 5, 695–702 (2008).

10. Tinevez, J.-Y. et al. TrackMate: An open and extensible platform for single-particle tracking. Methods 115, 80–90 (2017).

11. Beheiry, M. E. et al. A Primer on the Bayesian Approach to High-Density Single-Molecule Trajectories Analysis. Biophys J 110, 1209–15 (2016).

12. Masson, J.-B. et al. Mapping the energy and diffusion landscapes of membrane proteins at the cell surface using high-density single-molecule imaging and Bayesian inference: application to the multiscale dynamics of glycine receptors in the neuronal membrane. Biophys J 106, 74–83 (2014).

13. Calderon, C. Data-Driven Techniques for Detecting Dynamical State Changes in Noisily Measured 3D Single-Molecule Trajectories. Molecules 19, 18381–18398 (2014).

14. Calderon, C. P. & Bloom, K. Inferring Latent States and Refining Force Estimates via Hierarchical Dirichlet Process Modeling in Single Particle Tracking Experiments. Plos One 10, e0137633 (2015).

15. Buser, C. et al. Quantitative investigation of murine cytomegalovirus nucleocapsid interaction. J Microsc-oxford 228, 78–87 (2007).

16. Hoffman, D. P. et al. Correlative three-dimensional super-resolution and block-face electron microscopy of whole vitreously frozen cells. Science 367, (2020).

17. Xu, C. S. et al. Enhanced FIB-SEM systems for large-volume 3D imaging. Elife 6, e25916 (2017).

18. Xu, C. S. et al. An open-access volume electron microscopy atlas of whole cells and tissues. Nature 599, 147–151 (2021).

19. Lowe, D. G. Distinctive Image Features from Scale-Invariant Keypoints. Int J Comput Vision 60, 91–110 (2004).

20. Heinrich, L. et al. Whole-cell organelle segmentation in volume electron microscopy. Nature 599, 141–146 (2021).

