## Supplementary figures and images for "Motion of single molecular tethers reveals dynamic subdomains at ER-mitochondria contact sites"

### Supplemental Figures + Captions

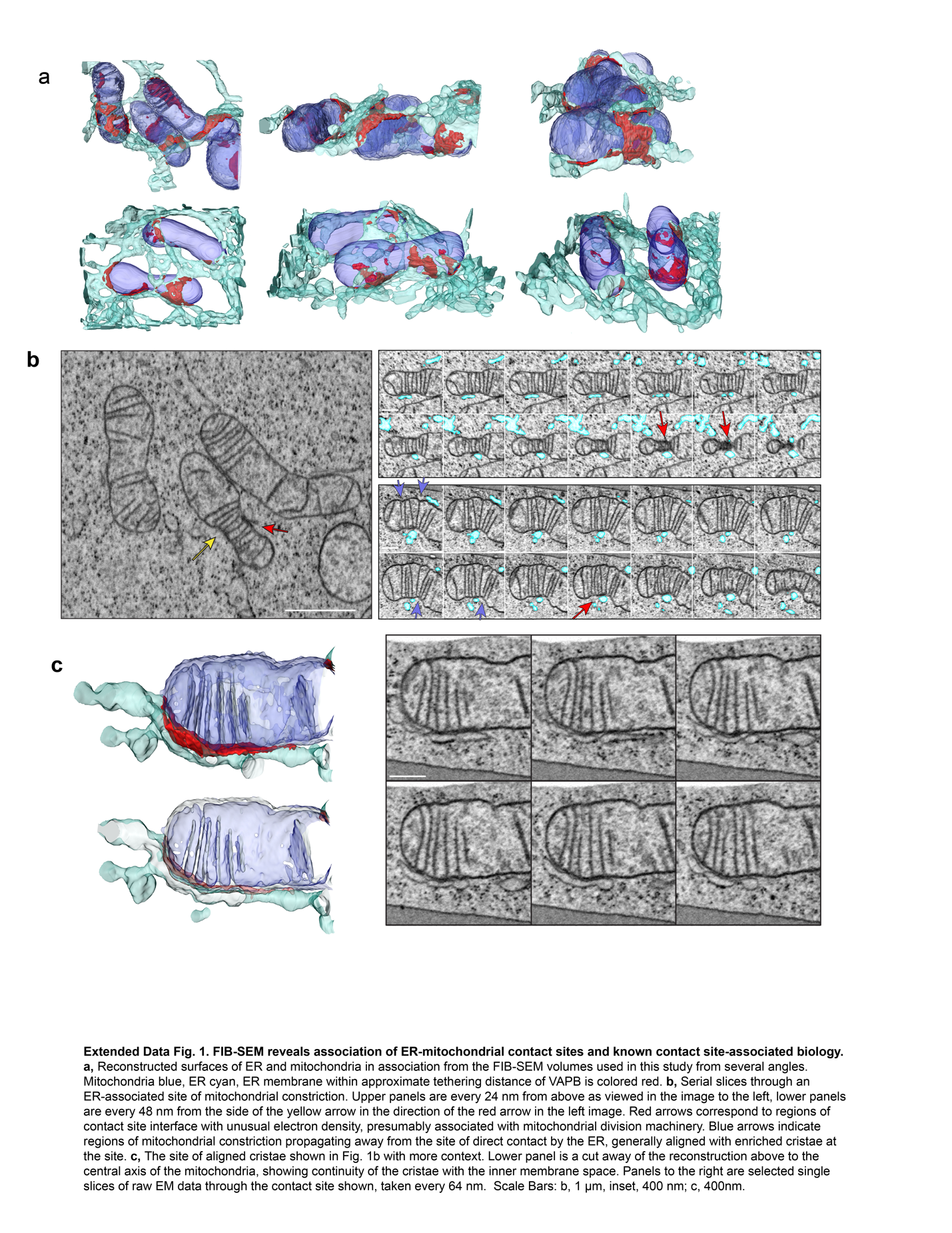

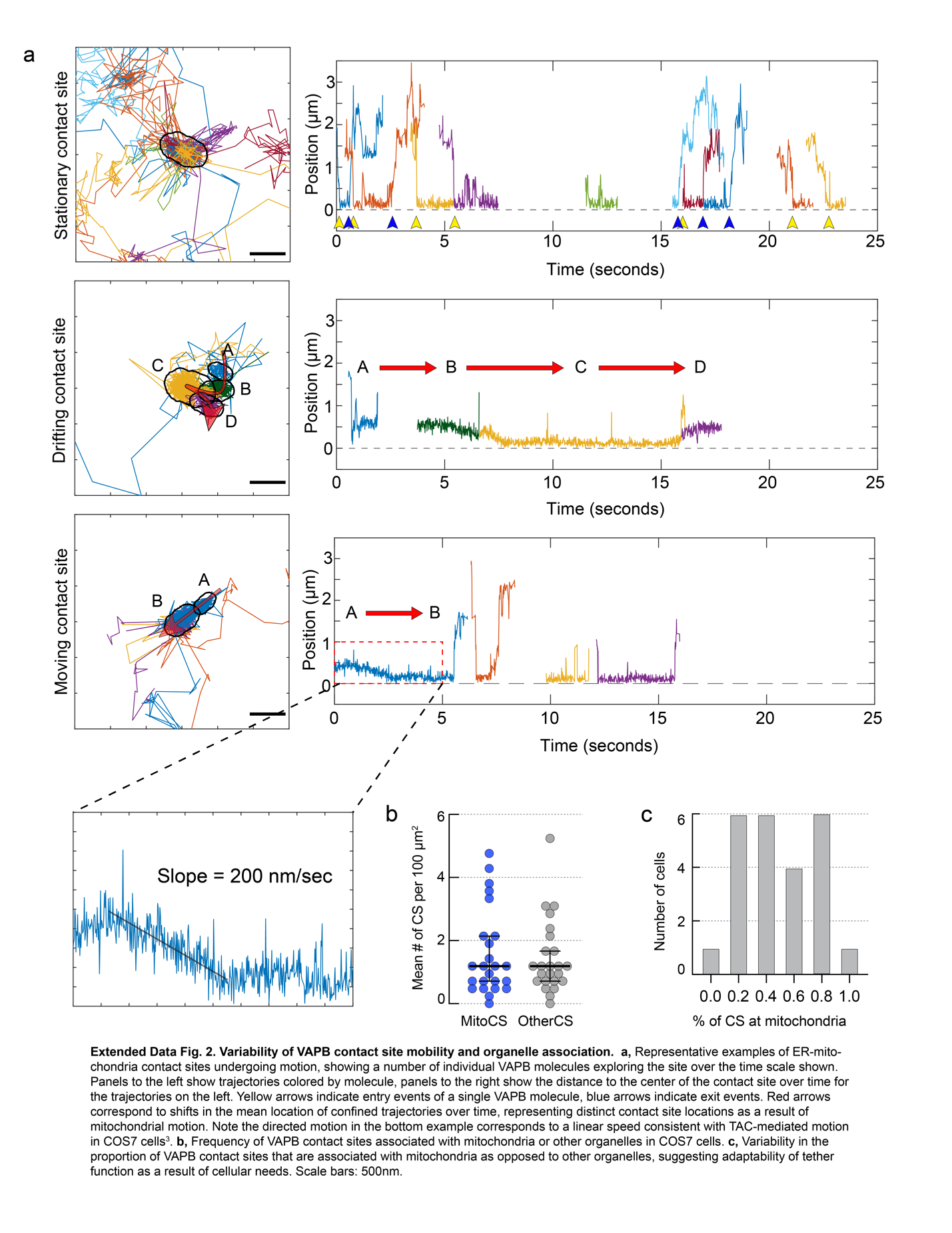

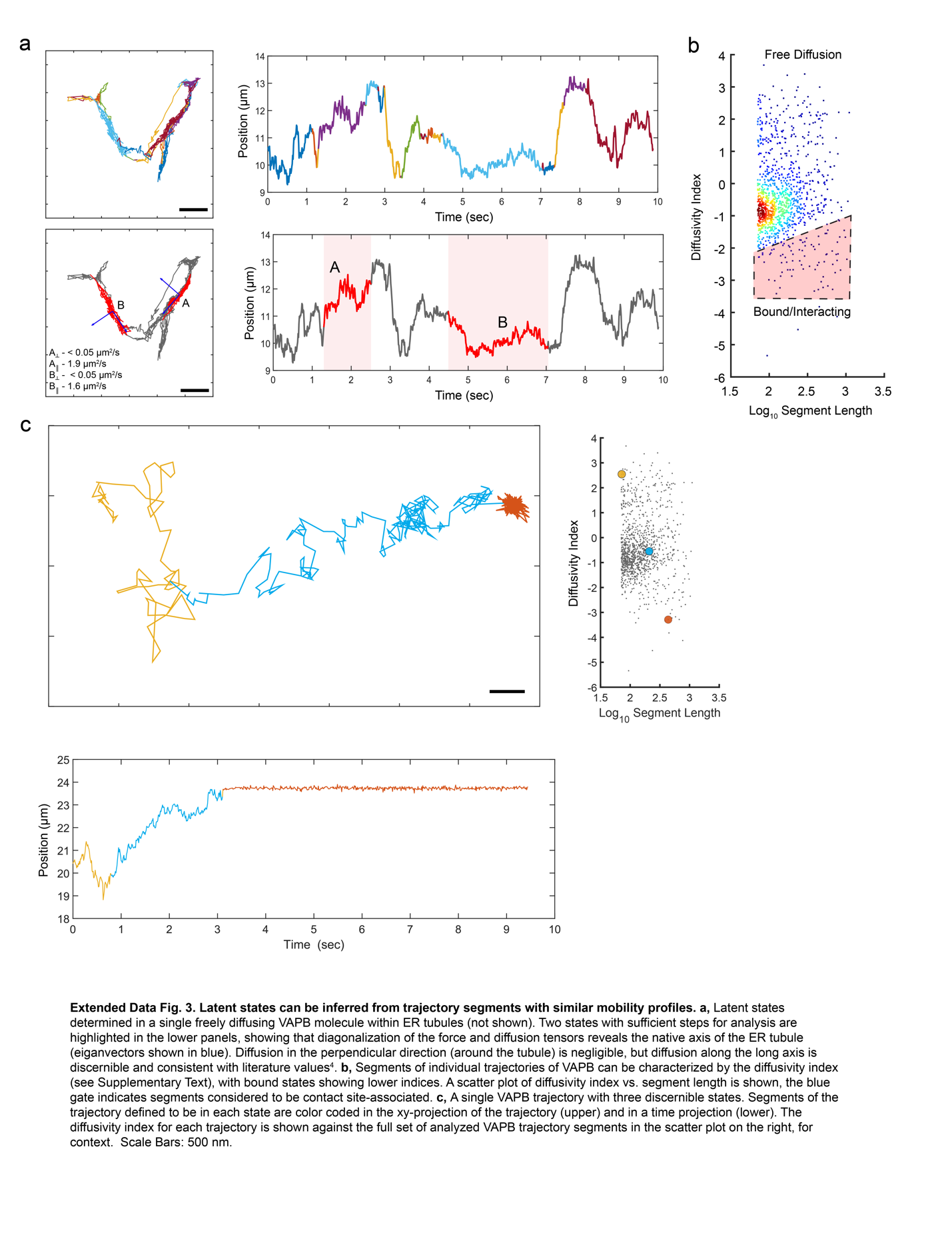

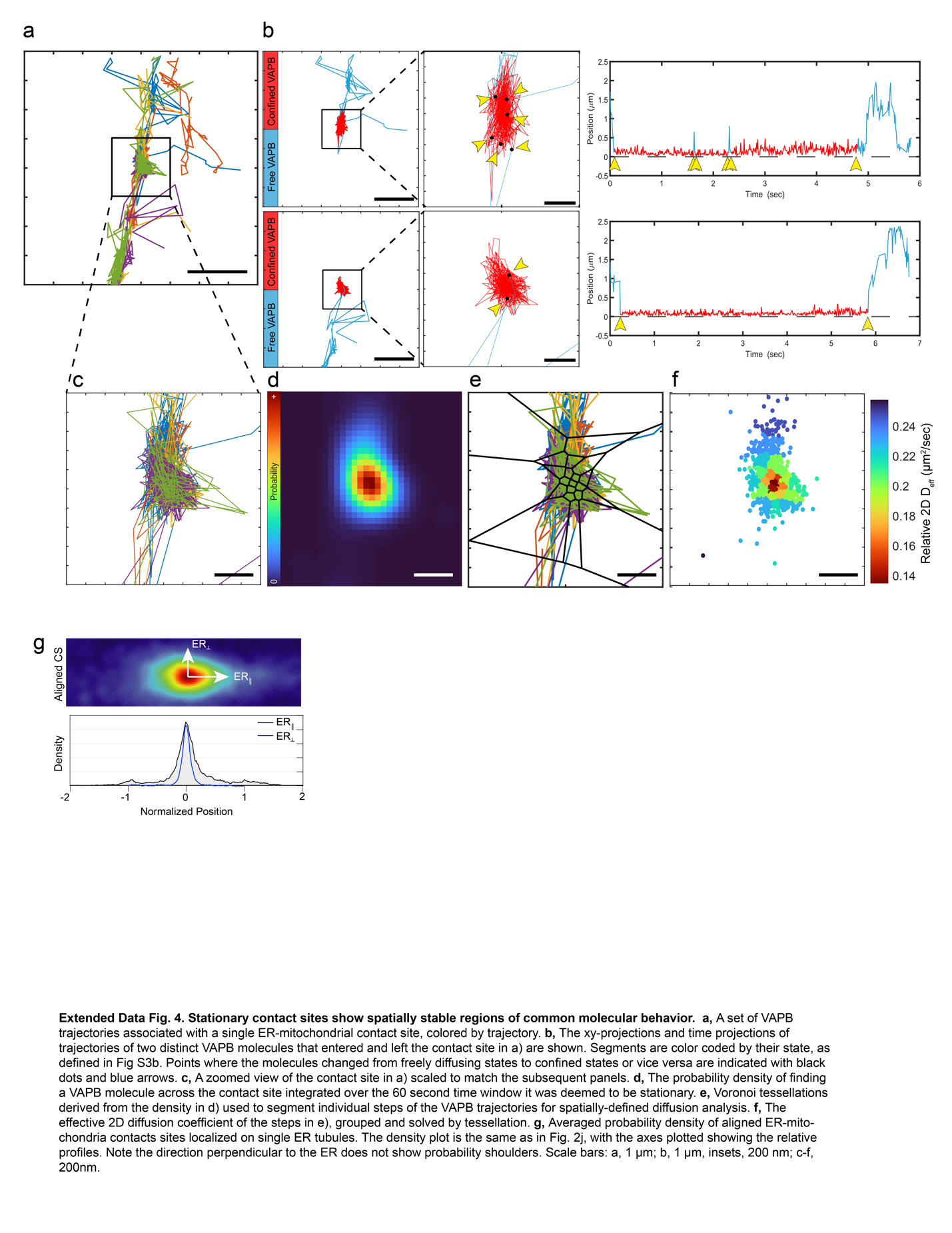

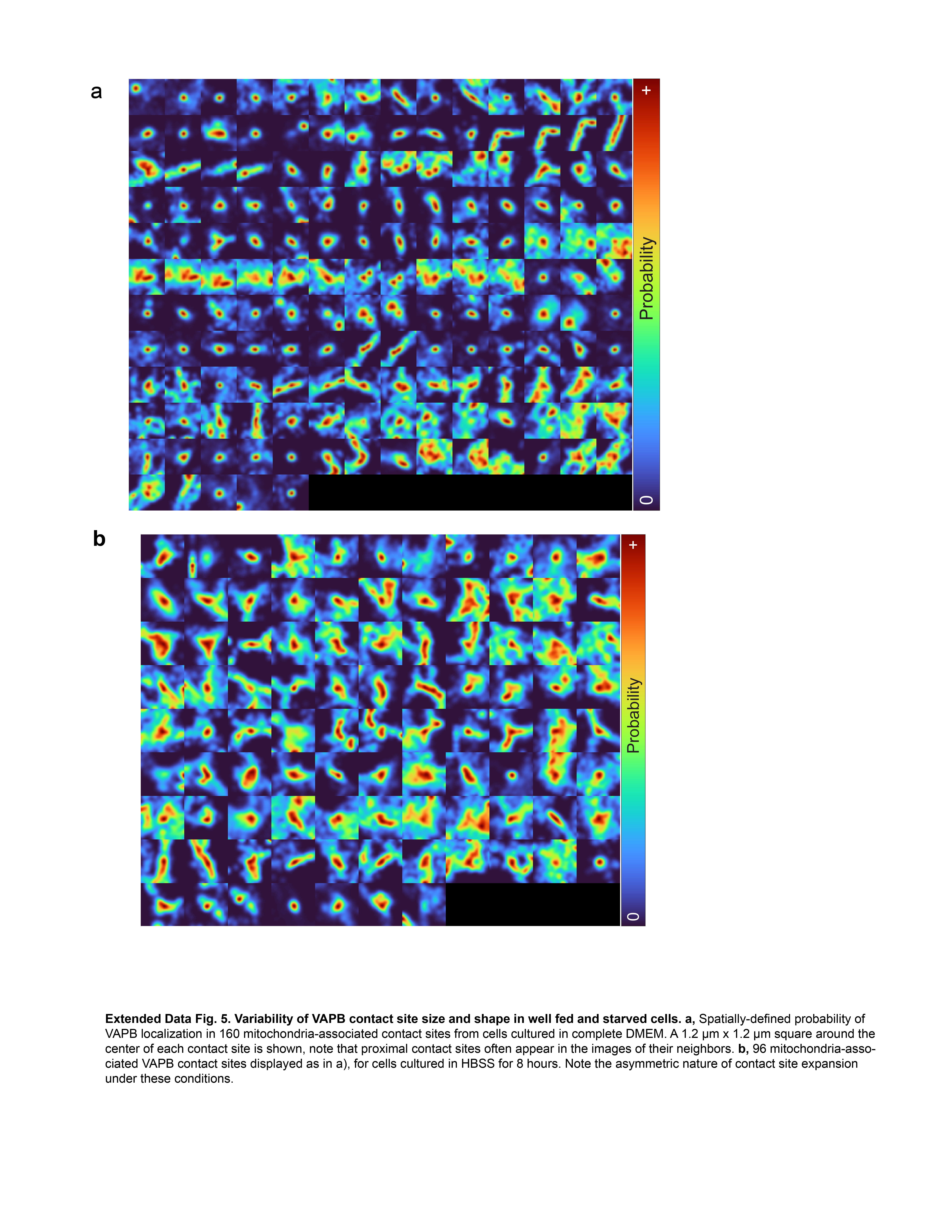
