## Supplementary Text and Methods for "Motion of single molecular tethers reveals dynamic subdomains at ER-mitochondria contact sites"

Supplemental Information

**TECHNICAL LIMITATIONS TO STUDYING CONTACT SITES BETWEEN ORGANELLES**

Contact sites between organelles have been extensively studied over the past forty years, primarily through reliance on traditional biochemistry and electron microscopy, though light microscopy has increasingly become an important validating tool in recent years^1,2^. These structures are particularly challenging to study, however, as they are generally very small, very dynamic, and very sensitive to experimental perturbation. We briefly describe here the risk of artifact associated with studying contact sites between organelles using existing technologies, which primarily fall into three categories. **Identification artifacts** are errors in the correct labeling of any potential organelle-organelle interaction as a specific contact site. **Perturbation artifacts** are errors in the measurement or conclusions drawn about contact sites as a result of the method of assaying them. **Averaging artifacts** are errors from the inherent loss of specificity that comes with pooling many independent samples together, either many cells with varied contact sites or even many diverse contact sites in a single cell. We detail sources of each of these errors we are aware of below.

**Sources of Identification artifacts**

1. Fixation—the number of apparent contact sites in EM data is significantly increased upon chemical fixation^3^, so clear establishment of what was a contact site in the unfixed cell is challenging. This may partially be the result of dehydration during EM sample preparation and thus may be avoided in fixed light microscopy experiments, but since this has not been clearly demonstrated to be the case, live cell experiments are preferrable.
2. Nonspecificity of EM labeling—in the absence of correlative optical microscopy, it is challenging to know that just because two membranes are within a certain distance that a specific contact has been established. Deviations from the expected shape, electron density, or associated structures can be evidence of specific contact, but still cannot be easily associated with specific machinery. This is underscored by the data in this paper (e.g.-Fig.2h-i), where regions of known contact sites within tethering distance were shown to avoid membrane deformation that is expected for regions of high-density tethering.
3. Nonspecificity of colocalization approaches—Use of colocalization in diffraction-limited microscopy has become a popular tool for identifying contact site location in live cells^4–6^. Given the limited spatial resolution of light microscopy techniques, many locations in the cell will have two organelle compartments within a voxel of one another that may not be directly interacting. Some groups have addressed this by simply counting sites where contact-associated functions occur^4,7^, but identification of *bona fide* contact sites where the desired function did not occur is then not possible. Others have required temporal associations^6^, reasoning that two organelles should not stay in proximity unless directly tethered. This is probably a good strategy at the periphery of tissue culture cells, where this lots of space for organelles to drift^8^, but is likely to create many false positives in the dense perinuclear area where mechanical crowding may hold organelles together even when not specifically tethered.
4. Sampling bias from low coverage in EM—EM-based approaches convey very high spatial resolution, but this elevated information density comes with an obligatory loss in throughput. A number of studies have shown that single slices through a cell as are common in transmission EM studies are not necessarily good quantitative representations of the full cell volume^8–13^, and the heterogeneity in contact sites described in this study (Extended Data Fig. 2b-c) suggest tens of cells must be sampled to capture the diversity of even a single tether ant contact sites. Thus, this tool must be coupled to higher throughput approaches like biochemistry and fluorescence microscopy.
5. Incomplete penetrance of specific labels—The limitations of colocalization-based approaches have led many groups to shift to using more specific labeling technologies like FRET sensors^14,15^, fluorescence complementation approaches^16,17^, and dimerization dependent fluorescent proteins (ddFPs)^18^. These significantly increase the specificity of labeling, since no signal is generated unless the two membranes are in sufficiently close proximity for complementation to occur. However, it is unclear the proportion of contact sites these labels can access. Presumably, they can access contact sites where the membranes are close enough for the synthetic tethers to reach one another, but contact sites in some contexts have been described to be much larger than the “conventional” values of 10-30nm, even in excess of 50 nm^19^—these complementation approaches may not access these interfaces. Additionally, existing protein or lipid complexes at contact sites may affect how easily synthetic tethers can access the interface.

**Sources of Perturbation artifacts**

1. Fixation—Performing the throughput of EM required to quantitatively measure the effects of fixation on contact site size, structure, membrane spacing, or composition has not been possible, but it seems likely some or all of these things could be affected^3,20,21^. Future work will be needed to determine the scale of this, but cryo-based approaches, though laborious, may be needed to accurately extract the nanoscale structure of contact sites.
2. Specific label-induced expansion—Given the incredible fluidity of the ERMCS interface described in this work, it seems likely that they will be very sensitive to perturbation even by relatively innocuous experimental protocols. For instance, the measured affinity of ddFPs is in a similar range to that of VAPB for FFAT-motifs^22^, and this is already a considerably lower stability interaction than complementation tools like more traditional spit fluorescent proteins that are commonly used. As we show in this work (Fig. 3), the size of the ER membrane necessitates that in most conditions the availability of the mitochondrial tether dominates the expansion or contraction of the contact site, and synthetic tethers by definition increase the availability of tethers in the mitochondria. Thus, these are likely to cause contact site expansion over time, even if the components are low affinity or of low temporal stability.
3. Specific label-induced protein displacement—Given our results with VAPB density in the central subdomain of the contact site, it seems likely that this extreme enrichment of specific tethers in this region may have an effect on the ability of other contact site proteins to access this space. Although we have not directly examined this, it seems likely synthetic tethering pairs would also be capable of this effect.
4. Specific label-induced membrane spacing artifacts— As even the low affinity interactions of VAPB and PTPIP51 can cause significant expansion of contact sites at defined membrane spacing^23–25^, it’s possible that the low affinity synthetic tethers may also have this effect.

**Sources and effects of averaging artifacts**

Even when only using VAPB as a probe, cells likely have at least hundreds of contact sites between the ER and the mitochondria. If there are ERMCSs in these cells that VAPB does not enter or mediate, it is likely this number is a significant underestimate of the total number of actively-tethered interfaces. Our data suggests significant cell-to-cell heterogeneity in contact site behavior even in the relatively simple system of immortalized cells in culture. Approaches that pool many diverse biological sites like biochemistry and even flow cytometry are likely to miss significant amounts of the subtlety in the system. Our data with P56S VAPB is a perfect example of this, where a very small fraction of the molecules are performing a medically-relevant behavior, but the vast majority of the molecules are not. This underscores the need for increasing the ease of experimental approaches with single biological event resolution.

**A note on the role of this work in solving some of these problems**

In this work, we have attempted to introduce tools and approaches that can be used to complement these existing experimental approaches in the literature. We do not yet know what proportion of contact sites detected by these means described above are mediated by VAPB, and it will be an exciting direction for future work to establish this and the role of other ER-mitochondria tethers at this interface. The sptPALM-based approach here provides an effective way to see all of the contact sites where VAPB is actively engaging in tethering and can in principle be combined with tracking of other tethers to generate a more complete vision of the heterogeneity and distribution of contact sites in single cells.

**FIB-SEM AND MEMBRANE RECONSTRUCTION**

**A note on reasoning for FIB-SEM and manual reconstruction approach**

High pressure freezing followed by freeze substitution and FIB-SEM has been successfully used to understand many of the structural aspects of organelle membranes in the absence of fixation-induced artifacts^3,8,10,21,26^, but the warping and staining effects of freeze substitution are still not well understood or mapped^21^. In principle, approaches performed in vitreous ice like cryo-ET, cryo-FIB-SEM, and cryo-EM avoid this issue, but in our hands these approaches do not offer sufficient contrast on ER membranes to perform nanoscale curvature or structural analysis. Additionally, the small volumes possible in these techniques make it hard to ensure that an ERMCS can be found within any given volume examined. As these technologies develop further, it will be exciting to examine this more closely, but current data suggests that warping effects in freeze substitution seem to effect local environments similarly, so the effects tend to be at the micron scale rather than the nanometer scale^21^. Thus, our local curvature analysis of the membrane is probably the best currently possible, but we will look forward to more careful analysis as cryo-based technologies for looking at membranes improve.

The most complete analyses of ER and mitochondria shape to date were also performed on high pressure frozen and freeze substituted FIB-SEM, but both used automated machine learning-based approaches for identifying the location and shape of membranes^8,19^. Although these provided a good sense of global ER and mitochondria shapes, we found that implementation of these types of approaches in our own data led to many errors in voxel classification in the tight space where the two opposing organelle membranes are in close proximity. Although these errors were often only a single voxel, they had dramatic effects on the curvature for the resulting triangulated surfaces, inhibiting our ability to analyze nanoscale membrane curvature in the contact sites themselves. Thus, we opted for a much smaller sample size and the somewhat laborious method of manual annotation of the data, but future work will examine the potential of training automated networks to minimize these types of artifacts.

**Method for smoothing and triangulation parameters**

The voxels in our FIB-SEM data are approximately 8nm in all dimensions, which is not sufficiently far from the scale of membrane curvature to avoid significant contributions to local curvature analysis. In order to minimize this source of error, we reasoned that we could use *a priori* information about the shape of the organelle membranes themselves. Significant effort in the field has been contributed to understanding the local shape of membranes, both at the level of high-resolution transmission EM and through modeling and prediction from *in vitro*-derived biophysical parameters. Thus, we chose smoothing parameters for each organelle surface that are selected to create a curvature that is most close to the established literature values. These parameters are implemented blindly to all the data in our volumes to avoid bias. All smoothing and triangulation steps were performed as an automated analysis pipeline in Amira that is available upon request.

**Method for curvature analysis**

Once triangulated surfaces were generated for the ER, we calculated the local curvature by fitting a quadratic form to the surface at each triangle using a 20-triangle neighborhood (approximately 25-30 nm, in our data). This process was iterated five times to smooth potential voxelation artifacts. The resulting eigenvectors and corresponding eigenvalues (C_1_ and C_2_) represent the principal curvature axes of the surface at this resolution, and we calculated the resulting mean curvature value directly at each triangle as:

$$Mean Curvature = \frac{C_{1}+C_{2}}{2}$$

Thus, convex regions have positive curvature, concave regions have negative curvature, and saddle points have net zero curvature. Note this is different from mean gaussian curvature that is often used in the literature, where saddle points would be defined as negative curvature and both convex and concave structures have positive Gaussian curvature.

**MOLECULAR BIOLOGY AND PLASMIDS**

**A note on construct choices**

Performing sptPALM in the ER requires an ER counterstain to inform the correct localization linkages that are needed to generate trajectories. We found that counterstains targeted to the membrane like mEmerald-Sec61b were good labels, but at high expression levels they showed a detectable effect on the diffusive properties of the single molecule tracers in the membrane, presumably as the result of molecular crowding. Thus, we elected to use an ensemble marker for the ER that resides entirely in the lumen to avoid perturbing tether motion.

Since VAPB also tethers FFAT-containing proteins on a number of non-mitochondrial membranes (reviewed here^27,28^), we required two labels (one for ER and one for mitochondria) in order to identify the contact sites that were associated with mitochondria (see Fig. 1j and Extended Data Fig. 2b and 2c). Although we could not directly visualize the mitochondrial tether and test its sensitivity to OMM labeling, we decided to target our mitochondrial tether to the mitochondrial matrix to avoid this potential.

The selection of two ensemble fluorescent labels that must be run simultaneously with sptPALM is challenging, since labels that are either too far red shifted or blue shifted can perturb the experiment. Fluorescence that is too red-shifted can create false localization or decrease true localization precision by elevating the background, and fluorochromes in this excitation profile are susceptible to bleaching by the 647nm laser used for single molecule localization. Fluorescent labels that require significant blue-shifted excitation light will cause too much photoconversion of PA-JF646 and lead to localizations that are too dense to unequivocally track. Unfortunately, in many cases these complications are not obvious during data acquisition, and only become clear after a laborious processing pipeline. To minimize the wasted effort this entails, we selected the two ensemble markers used throughout this paper that gave us the largest proportion of usable data (PrSS-mEmerald-KDEL and mitoRFP), though these two markers can induce ER stress at late time points. As a result, all data presented in this paper is collected at less than 24 hours post transfection, where the ER morphology and function were deemed to still be reasonably healthy (and the motion of single molecule tracers was consistent with earlier time points).

**Cloning strategy and construct design**

The ER ensemble marker (PrSS-mEmerald-KDEL) was generated because in our hands mEmerald showed the best balance between the required brightness, obligate photostability, and toxicity effects of overexpression of the green fluorophores. Briefly, PrSS-mEmerald-KDEL was generated by replacing the mRFP cassette in ER-mRFP using NEBuilder to exchange the catalytic cores of the fluorescent protein with homologous arms. The plasmid was sequence verified before use, and the plasmid sequence and map are available at Addgene or by request.

HaloTag-TA was generated by PCR amplifying a codon optimized version of HaloTag^29,30^ and adding flanking AgeI and BsrG1 restriction sites using the primers in table S1. The resulting product was inserted into mEmerald-Sec61b-C1 by digesting both constructs with AgeI and BsrG1 and ligating the purified products. The plasmid was sequence verified before use and the map and sequence are available at Addgene or by request.

All other single molecule tracer constructs were made from HaloTag-N1 and HaloTag-C1 backbones, which were generated by replacing the core of EGFP-N1 and EGFP-C1 with the same HaloTag core as used for HaloTag-TA above. This was performed using NEBuilder and the primers described in table S1 according to the manufacturer’s recommendations. Note the very C and N terminus of EGFP were retained where the fused to the linker sequence, to avoid folding issues and increase linker flexibility, since the GFP-tagged versions of the proteins used in this paper have been previously shown to be functional^31^.

To ensure that the HaloTag-linked versions of VAPB still retained their contact site-associated interactions, we generated both N- and C- terminally linked versions of VAPB by inserting the full length VAPB (amino acids 1-243) into both HaloTag-N1 and HaloTag-C1, using NEBuilder. The resulting constructs were sequence verified and then checked for behavior with sptPALM. Both constructs showed clear interactions at the interface of ER and mitochondria, but a significant number of the C-terminally tagged VAPB molecules were freely diffusing in three dimensions within the cytoplasm. We presumed this represented a defect in the tail anchor insertion pathway as a result of adding the bulky HaloTag to the C-terminus, which must be passed through the membrane and face the ER lumen. As a result, we performed the remainder of the experiments in the paper using the N-terminally tagged versions of VAPB. The maps and plasmid sequences are available at Addgene or by request.

The VAPB construct with the N-terminus deleted was generated by inserting the last 32 amino acids of VAPB (amino acids 212-243) into HaloTag-C1 using NEBuilder. This deletion removes both the PTPIP51-interacting domain and the dimerization domain, essentially leaving only the transmembrane tail anchor and short sequence on either side. This proved to be sufficient for targeting to the ER, but it did not convey any detectable specificity to the motion of the tethers in the contact sites themselves (Fig. 1k-m), suggesting that VAPB does not use it’s TM domain to sense the unique lipid environments of the contact site. An annotated map and plasmid sequence are available at Addgene or by request.

The VAPB-N-terminus fused to the tail anchor control from Sec61b was generated by PCR amplifying the synthetic HaloTag-VAPB fusion from HaloTag-VAPB, but only amplifying the N-terminal region (amino acids 1-211). This construct was inserted in place of mEmerald in mEmerald-Sec61b-C1 using NEBuilder. The resulting construct was sequence verified before use, and the map and sequence are available at Addgene or by request. This construct was also entirely targeted to the ER membrane, but it did show distinct regions of interaction at the mitochondria-associated contact sites (Fig. 1k-m).

HA-PTPIP51 has been previously shown to retain its function^25^, but the protein’s ability to tolerate a larger tag was unclear. However, we wished to perform experiments in live cells, where the HA-tag was not a reasonable labeling option. In order to surmount this, we extracted the full length PTPIP51 (amino acids 1-470) from HA-PTPIP51 using the primers in table S1 and fused it directly to the modified EMCV IRES sequence from pHAGE-Tet-STEMCCA and mTagBFP2 from mTagBFP2-N1 using NEBuilder. The full sequence was inserted into EGFP-N1 replacing the EGFP. The resulting plasmid map and sequence available at Addgene or by request. The construct was sequence verified before use and allowed the overexpression of PTPIP51 without any additional tags, while simultaneously producing mTagBFP2 in the cytosol to mark the transfected cells.

**IMAGING STRATEGY AND IMAGE PREPARATION/PROCESSING**

**A note on imaging strategy, laser power, and image preparation**

One of the most challenging aspects of sptPALM with multiple simultaneously-collected ensemble labels is that green labels that are often preferred for imaging require 488nm light for excitation. 488nm light can inefficiently cause photoactivation of the PA-JF646 dye used for sptPALM in our experiments. We tested a number of conditions and constructs, but found that even without any 405 radiation to photoconvert the dye, nonspecific photoactivation by the 488nm laser became problematic for us if more 100µW were applied to the back aperture (this corresponds to only approximately 272 mW/cm^2^). Thus, if cells could not be visualized beneath this power, they were not able to be used for ER-based sptPALM. This is generally not enough excitation light to visualize even the brightest green ER structural labels when imaging camera exposure times are less than 20 msec, so we required some image processing steps to improve the signal to noise in the structural channels, described below.

**Imaging speed requirements for ER motion**

We have previously shown that even at 95Hz some molecular motion of membrane proteins in under sampled^32^, and localization distortion from molecular motion is present even at this speed^33^. Keeping molecular steps small enough to identify single ER junction transitions requires a minimum of 15-20Hz in the peripheral ER^32^. However, the structure itself does not require such fast imaging to track unequivocally. A number of papers throughout the years have quantified the high-speed motion of the peripheral ER using fluorescent labels for the structure^9,34–44^. Major structural rearrangements like tubule extensions^34–36^, ring closures^45^, and three-way junction formation^46–50^ occur on a time scale of hundreds of milliseconds to seconds. We and others have described very high frequency oscillations in ER tubules that occur in excess of 40-50Hz as a result of cytoplasmic fluctuations^9,44^, but these are often subdiffraction-limited in size and occur perpendicular to the central axis of ER tubules (and as such do not confound single molecule linkages along the tubules). Thus, we reasoned that if our effective imaging speed for the ER structure was in the range of 10Hz, we should still achieve Nyquist sampling for structural motion even if the structure was not imaged as quickly as the single molecule localizations required.

**Image filtering and preparation**

To accomplish this, the ER and mitochondrial channels were subjected to a 10-frame median filter in the time domain before being utilized for subsequent tracking support (effective exposure time, 110msec). The single molecule channel was left unprocessed, to avoid distorting the localizations or confusing the trajectories. The resulting filtered images still occasionally showed signs of bleaching in one or both channels, so we also performed a simple ratio bleach correction before subsequent image processing. The temporal median-filtered and bleach corrected image data is used for all ER and mitochondria images in this paper.

The curation of trajectories that is required to avoid linkage artifacts in sptPALM of ER proteins is a relatively laborious process as it was performed in this study. In order to make this as simple as possible, we performed a number of additional steps to help the trajectory curation. First, we generated a very crude mask of the ER using a simple threshold to generate a binary image and then performed a single-pixel dilation three times sequentially. Since the pixels are very large (160nm), this functionally expanded the mask to include all regions of the cell that contained ER. Localizations that fell outside this mask we removed before performing tracking. These events were rare, likely representing unbound dye, but in practice they can be distracting while setting tracking parameters.

**SINGLE MOLECULE TRACKING AND TRAJECTORY ANALYSIS**

**A note on complications of tracking in arbitrary organelle geometries**

Single particle tracking (SPT) has proven to be a powerful tool for the analysis of biological phenomena at the molecular level in diverse environments^51–53^. Due to the high precision nature of the technique, nearly every step in the experiment and analysis pipeline from localization to tracking to analysis makes relatively heavy use of mathematical approaches. It is important to note that most of these approaches make some inherent assumptions about the boundary conditions and symmetry of the system. When tracking in contexts where symmetry is reasonably uniform (such as the ventral plasma membrane of a cell cultured on glass) these assumptions introduce minimal error into the analysis. However, when localizing and tracking molecules in the complex environment of the morphologically complex cytoplasmic organelles, these assumptions can lead to significant artifacts. In this section, we briefly discuss sources of error and controls that can be run to minimize them or account for them that we have performed throughout this study.

**Effects of organelle structure on linkage artifacts and trajectory formation**

Most traditional linking algorithms for SPT use an initial linking step based on nearest neighbor analysis that is then subjected to a subsequent optimization function^51,54^. In the context of an arbitrarily shaped organelle like the ER or the mitochondria, the nearest localization in 2D space is often not the nearest neighbor in the space in which the particles reside (and in the case of the mitochondria is often not even in the same organelle). We found by manual inspection of the data that even at modest linking densities the majority of trajectories showed steps that crossed at best prohibitively large distances through the structure, if not transitions that should be altogether forbidden. Future work may develop automated algorithms to use the underlying structure to directly inform the linking process, but in this study, we performed this step manually. The experimentalist was blinded to the condition and went through each frame of the time lapse manually breaking trajectory linkages that stepped over the polygonal spaces of the ER, which would have required a molecule to travel much more quickly through a circuitous route than is reasonable. As a qualitative validation, this process was performed on datasets collected in similar ER structures at a range of photoconversion densities, and subsequent work was performed at densities where the linkage properties were insensitive to localization density.

**Effects of organelle structure on ensemble analysis of trajectories**

Most statistical approaches for analyzing single particle trajectories make some base assumptions about the symmetry or at least isotropy of the system, especially the contribution of thermal fluctuations and Brownian motion^51^. Molecules confined to organelles without a high degree of symmetry are not necessarily subject to these assumptions, since diffusion can make significant contributions to apparently “directional” motion if some dimensions are more constrained than others. We have previously shown that both traditional mean-squared displacement (MSD) analysis and velocity autocorrelation produce erroneous results in the endoplasmic reticulum if the organelle shape is not accounted for^32^. This approach can be used effectively in ER tubules, where *a priori* knowledge of the membrane shape allows the effective reduction to a one-dimensional problem, but without an orthogonal method of assaying contact site structure at this resolution it cannot be used to understand particle motion in the contact site. Thus, in this study we utilized a simple approach to minimize the error rather than try to remove it completely.

Briefly, we reasoned that since molecular diffusion operates over much faster time scales than organelle restructuring, in the limit of vanishingly small subregions of the ERMCS, most molecules within a reasonable time window should see approximately the same structure (Extended Data Fig. 4). As a result, we make the assumption that they should experience approximately similar diffusion and energetic environments, treating them as a constants and solving the overdamped limit of the Langevin equation for the trajectories in each neighborhood:

$$\frac{d\overset{⃑}{r}}{dt}=\frac{\overset{⃑}{F}(\overset{⃑}{r})}{\gamma(\overset{⃑}{r})}+\sqrt{2D(\overset{⃑}{r})}\xi(t)$$

where the left term is a generic “drift” term that is left to account for nondiffusive aspects of the motion, and the right term estimates the Brownian motion in the neighborhood^55^. Additionally, the contribution of the “drift” term used to solve the equation is deliberately unconstrained to avoid bias about the nature of contact site binding, and in this region where confinement to the site is significant it does contribute. However, this contribution serves to decrease the signal to noise for changes in D_eff_ across the contact site, and such biases against our conclusions of subdomain-specific variation. This approach of breaking diffusion landscapes down into smaller subdomains has been used effectively in a variety of biological structures^55–57^, though we caution that this does not completely remove the effects of structural variation, it simply reduces them. As a result, the numbers and D_eff_ extracted using this technique are not directly comparable to those collected using the more precise approaches based on single trajectories detailed below or our previously published work^32^, which are in close agreement. However, this approach has served as an effective way to map the landscape of contact sites qualitatively, since molecules within the same contact site are likely subject to similar environments and analysis of hundreds of contact sites provided very consistent results.

One weakness of this approach, however, is that it not very effective in parsing changing state behavior, especially when the states are not uniformly changing as a function of location. Thus, in healthy ERMCSs, molecular behavior is more or less consistent within each neighborhood and low D_eff_ regions correspond well to where molecules are more likely to be located (Fig. 2, Extended Data Fig.4). However, in the P56S VAPB tracking, subdomains that trap P56S VAPB can often also have normally diffusing molecules present. Thus, these present as domains of very low diffusion using this approach, but future work will be needed to ascertain what proportion of this motion is truly “slowed diffusion” as opposed to increasing confinement as a result of lateral aggregation to other VAPB molecules or more stable binding to PTPIP51. There is some evidence for both of these as potential options^25,58–60^ (despite ensemble biochemistry suggesting a loss of P56S interaction with PTPIP51^58,61–63^, probably the result of the majority being in ER-localized aggregates^64^), but parsing these states will require a more complex approach (see below) beyond the capacity of our current technology.

**Effects of organelle structure on single trajectory analysis and latent state determination**

Ensemble-averaged approaches like MSD or velocity autocorrelation have significant limitations in real biological systems in that they by necessity have trouble with temporally or spatially varying forces or contributions to motion^65^. In a dynamic and complex organelle structure like the ER or mitochondria, both spatial and temporal variation in the organelle’s shape have significant contributions to molecular motion^32^. Approaches like traditional Hidden Markov Models can surmount this problem, but they require two significant constraints that are in practice nearly impossible to achieve with such asymmetric boundary conditions^66–68^. First, trajectories must be long enough to allow statistical convergence of the model, which can in practice be challenging to achieve with single molecule probes due to natural bleaching of the fluorochrome under the requisite levels of radiation. Second, they require an accurate upper bound on the number of states possible in the system. In practice, this requirement is essentially impossible to achieve in the highly complex and variable structure of the ER, since variations on local curvature, topology, connectivity, and protein/lipid content can produce many functionally distinct states. In this work, we take advantage of the superior photostability of new photoconvertible fluorochromes^69–71^ to address the first point, and a model-independent way of determining latent states to address the second^72,73^.

The complete approach used in this work is detailed elsewhere^72,73^, but the resulting segments were often broken such that they correctly identified structural differences in the underlying ER (Extended Data Fig.3a). For each segment of a trajectory that was long enough (generally >150 steps), an implementation of the two-dimensional overdamped Langevin equation was solved:

$$d{\overset{⃑}{\boldsymbol{r}}}_{t} = \gamma^{-1}\overset{⃑}{\boldsymbol{F}}({\overset{⃑}{r}}_{t} )dt + \sqrt{2D}d{\overset{⃑}{\boldsymbol{B}}}_{t}$$

(Note that although the model and lettering notation are the same as in the section above, these values do not directly correspond to one another and are calculated along different dimensional spaces). The kinetic values of ***D_eff_*** and ***F*** along each eigenvector of the two component matrices was estimated using traditional maximum likelihood estimation. As a sanity check, we noted that most isolated segments of VAPB diffusion in ER tubules were correctly broken into axes that were parallel or perpendicular to the central cylindrical axis of the ER tubule, and the contribution of diffusion to the perpendicular axis was negligible. Additionally, we noticed that the diffusion of VAPB along the tubule axes simplified to values that are nearly identical to those we have estimated for ER tubule diffusion of another tail anchored protein by a completely unrelated mathematical approach^32^.

**Use of Diffusivity Index to classify latent states**

Since we do not constrain the relative contributions of thermal motion and external forces within each segment, the relative contribution of these two terms provides a rough measure of how closely a particular trajectory segment is behaving to unconstrained diffusion. Thus, we introduce a simple “Diffusivity Index” (Extended Data Fig. 3b) for each segment of sufficient length which is defined as the ratio of the thermal term to the force term:

$$i.e., Diffusivity Index \approx\sum_{Traj. Seg.} \frac{\sqrt{2D}}{\gamma^{-1}\overset{⃑}{\boldsymbol{F}}\left( {\overset{⃑}{r}}_{t} \right)}$$

VAPB trajectories in complex ER structures like ER matrices or sheets with two dimensions of freedom show a high Diffusivity Index (though not necessarily a high D_eff_), while trajectories primarily localized in isolated tubules or thin structures tend to show medium diffusivity indices (yellow and blue, Extended Data Fig. 3c). Trajectories confined to contact sites or immobilized through binding or aggregation show very low diffusivity indices (orange, Extended Data Fig. 3c).

**Extended Data Fig. 1. FIB-SEM reveals association of ER-mitochondrial contact sites and known contact site-associated biology. a,** Reconstructed surfaces of ER and mitochondria in association from the FIB-SEM volumes used in this study from several angles. Mitochondria blue, ER cyan, ER membrane within approximate tethering distance of VAPB is colored red. **b,** Serial slices through an ER-associated site of mitochondrial constriction. Upper panels are every 24 nm from above as viewed in the image to the left, lower panels are every 48 nm from the side of the yellow arrow in the direction of the red arrow in the left image. Red arrows correspond to regions of contact site interface with unusual electron density, presumably associated with mitochondrial division machinery. Blue arrows indicate regions of mitochondrial constriction propagating away from the site of direct contact by the ER, generally aligned with enriched cristae at the site. **c,** The site of aligned cristae shown in Fig. 1b with more context. Lower panel is a cut away of the reconstruction above to the central axis of the mitochondria, showing continuity of the cristae with the inner membrane space. Panels to the right are selected single slices of raw EM data through the contact site shown, taken every 64 nm. Scale Bars: b, 1 µm, inset, 400 nm; c, 400nm.

**Extended Data Fig. 2. Variability of VAPB contact site mobility and organelle association. a,** Representative examples of ER-mitochondria contact sites undergoing motion, showing a number of individual VAPB molecules exploring the site over the time scale shown. Panels to the left show trajectories colored by molecule, panels to the right show the distance to the center of the contact site over time for the trajectories on the left. Yellow arrows indicate entry events of a single VAPB molecule, blue arrows indicate exit events. Red arrows correspond to shifts in the mean location of confined trajectories over time, representing distinct contact site locations as a result of mitochondrial motion. Note the directed motion in the bottom example corresponds to a linear speed consistent with TAC-mediated motion in COS7 cells**^44^**. **b,** Frequency of VAPB contact sites associated with mitochondria or other organelles in COS7 cells. **c,** Variability in the proportion of VAPB contact sites that are associated with mitochondria as opposed to other organelles, suggesting adaptability of tether function as a result of cellular needs. Scale bars: 500nm.

**Extended Data Fig. 3. Latent states can be inferred from trajectory segments with similar mobility profiles. a,** Latent states determined in a single freely diffusing VAPB molecule within ER tubules (not shown). Two states with sufficient steps for analysis are highlighted in the lower panels, showing that diagonalization of the force and diffusion tensors reveals the native axis of the ER tubule (eiganvectors shown in blue). Diffusion in the perpendicular direction (around the tubule) is negligible, but diffusion along the long axis is discernible and consistent with literature values^32^. **b,** Segments of individual trajectories of VAPB can be characterized by the diffusivity index (see ), with bound states showing lower indices. A scatter plot of diffusivity index vs. segment length is shown, the blue gate indicates segments considered to be contact site-associated. **c,** A single VAPB trajectory with three discernible states. Segments of the trajectory defined to be in each state are color coded in the xy-projection of the trajectory (upper) and in a time projection (lower). The diffusivity index for each trajectory is shown against the full set of analyzed VAPB trajectory segments in the scatter plot on the right, for context. Scale Bars: 500 nm.

**Extended Data Fig. 4. Stationary contact sites show spatially stable regions of common molecular behavior. a,** A set of VAPB trajectories associated with a single ER-mitochondrial contact site, colored by trajectory. **b,** The xy-projections and time projections of trajectories of two distinct VAPB molecules that entered and left the contact site in a) are shown. Segments are color coded by their state, as defined in Fig S3b. Points where the molecules changed from freely diffusing states to confined states or vice versa are indicated with black dots and blue arrows. **c,** A zoomed view of the contact site in a) scaled to match the subsequent panels. **d,** The probability density of finding a VAPB molecule across the contact site integrated over the 60 second time window it was deemed to be stationary. **e,** Voronoi tessellations derived from the density in d) used to segment individual steps of the VAPB trajectories for spatially-defined diffusion analysis. **f,** The effective 2D diffusion coefficient of the steps in e), grouped and solved by tessellation. Scale bars: a, 1 µm; b, 1 µm, insets, 200 nm; c-f, 200nm.

**Extended Data Fig. 5. Variability of VAPB contact site size and shape in well fed and starved cells. a,** Spatially-defined probability of VAPB localization in 160 mitochondria-associated contact sites from cells cultured in complete DMEM. A 1.2 µm x 1.2 µm square around the center of each contact site is shown, note that proximal contact sites often appear in the images of their neighbors. **b,** 96 mitochondria-associated VAPB contact sites displayed as in a), for cells cultured in HBSS for 8 hours. Note the asymmetric nature of contact site expansion under these conditions.

1. Prinz, W. A., Toulmay, A. & Balla, T. The functional universe of membrane contact sites. *Nat Rev Mol Cell Bio* 21, 7–24 (2019).

2. Phillips, M. J. & Voeltz, G. K. Structure and function of ER membrane contact sites with other organelles. *Nat Rev Mol Cell Bio* 17, 69–82 (2016).

3. Pezzati, R., Bossi, M., Podini, P., Meldolesi, J. & Grohovaz, F. High-resolution calcium mapping of the endoplasmic reticulum-Golgi-exocytic membrane system. Electron energy loss imaging analysis of quick frozen-freeze dried PC12 cells. *Mol Biol Cell* 8, 1501–1512 (1997).

4. Friedman, J. R. *et al.* ER Tubules Mark Sites of Mitochondrial Division. *Science* 334, 358–362 (2011).

5. Lewis, S. C., Uchiyama, L. F. & Nunnari, J. ER-mitochondria contacts couple mtDNA synthesis with mitochondrial division in human cells. *Science* 353, aaf5549 (2016).

6. Valm, A. M. *et al.* Applying systems-level spectral imaging and analysis to reveal the organelle interactome. *Nature* 546, 162–167 (2016).

7. Hoyer, M. J. *et al.* A Novel Class of ER Membrane Proteins Regulates ER-Associated Endosome Fission. *Cell* 175, 254-265.e14 (2018).

8. Heinrich, L. *et al.* Whole-cell organelle segmentation in volume electron microscopy. *Nature* 599, 141–146 (2021).

9. Nixon-Abell, J. *et al.* Increased spatiotemporal resolution reveals highly dynamic dense tubular matrices in the peripheral ER. *Science* 354, aaf3928–aaf3928 (2016).

10. Weigel, A. V. *et al.* ER-to-Golgi protein delivery through an interwoven, tubular network extending from ER. *Cell* 184, 2412-2429.e16 (2021).

11. Heymann, J. A. W. *et al.* 3D Imaging of mammalian cells with ion-abrasion scanning electron microscopy. *J Struct Biol* 166, 1–7 (2009).

12. Felts, R. L. *et al.* 3D visualization of HIV transfer at the virological synapse between dendritic cells and T cells. *Proc National Acad Sci* 107, 13336–13341 (2010).

13. Bennett, A. E. *et al.* Ion-Abrasion Scanning Electron Microscopy Reveals Surface-Connected Tubular Conduits in HIV-Infected Macrophages. *Plos Pathog* 5, e1000591 (2009).

14. Csordás, G. *et al.* Imaging Interorganelle Contacts and Local Calcium Dynamics at the ER-Mitochondrial Interface. *Mol Cell* 39, 121–132 (2010).

15. Naon, D. *et al.* Critical reappraisal confirms that Mitofusin 2 is an endoplasmic reticulum–mitochondria tether. *Proc National Acad Sci* 113, 11249–11254 (2016).

16. Wilson, E. L. & Metzakopian, E. ER-mitochondria contact sites in neurodegeneration: genetic screening approaches to investigate novel disease mechanisms. *Cell Death Differ* 28, 1804–1821 (2021).

17. Harmon, M., Larkman, P., Hardingham, G., Jackson, M. & Skehel, P. A Bi-fluorescence complementation system to detect associations between the Endoplasmic reticulum and mitochondria. *Sci Rep-uk* 7, 17467 (2017).

18. Lee, J. E., Cathey, P. I., Wu, H., Parker, R. & Voeltz, G. K. Endoplasmic reticulum contact sites regulate the dynamics of membraneless organelles. *Science* 367, (2020).

19. Parlakgül, G. *et al.* Regulation of liver subcellular architecture controls metabolic homeostasis. *Nature* 603, 736–742 (2022).

20. Hicks, M. L., Brilliant, J. D. & Foreman, D. W. Electron Microscope Comparison of Freeze-Substitution and Conventional Chemical Fixation of Undecalificied Human Dentin. *J Dent Res* 55, 400–410 (1976).

21. Hoffman, D. P. *et al.* Correlative three-dimensional super-resolution and block-face electron microscopy of whole vitreously frozen cells. *Science* 367, (2020).

22. Ding, Y. *et al.* Ratiometric biosensors based on dimerization-dependent fluorescent protein exchange. *Nat Methods* 12, 195–198 (2015).

23. Stoica, R. *et al.* ALS/FTD‐associated FUS activates GSK‐3β to disrupt the VAPB–PTPIP51 interaction and ER–mitochondria associations. *Embo Rep* 17, 1326–1342 (2016).

24. Stoica, R. *et al.* ER–mitochondria associations are regulated by the VAPB–PTPIP51 interaction and are disrupted by ALS/FTD-associated TDP-43. *Nat Commun* 5, 3996 (2014).

25. Vos, K. J. D. *et al.* VAPB interacts with the mitochondrial protein PTPIP51 to regulate calcium homeostasis. *Hum Mol Genet* 21, 1299–1311 (2012).

26. Xu, C. S. *et al.* An open-access volume electron microscopy atlas of whole cells and tissues. *Nature* 599, 147–151 (2021).

27. Murphy, S. E. & Levine, T. P. VAP, a Versatile Access Point for the Endoplasmic Reticulum: Review and analysis of FFAT-like motifs in the VAPome. *Biochim Biophys Acta* 1861, 952–61 (2016).

28. Kors, S., Costello, J. L. & Schrader, M. VAP Proteins – From Organelle Tethers to Pathogenic Host Interactors and Their Role in Neuronal Disease. *Frontiers Cell Dev Biology* 10, 895856 (2022).

29. Los, G. V. *et al.* HaloTag: A Novel Protein Labeling Technology for Cell Imaging and Protein Analysis. *Acs Chem Biol* 3, 373–382 (2008).

30. Encell, L. P. *et al.* Development of a Dehalogenase-Based Protein Fusion Tag Capable of Rapid, Selective and Covalent Attachment to Customizable Ligands. *Curr Chem Genom* 6, 55–71 (2012).

31. Dong, R. *et al.* Endosome-ER Contacts Control Actin Nucleation and Retromer Function through VAP-Dependent Regulation of PI4P. *Cell* 166, 408–423 (2016).

32. Sun, Y. *et al.* Unraveling trajectories of diffusive particles on networks. *Phys Rev Res* 4, 023182 (2022).

33. Calderon, C. P. Motion blur filtering: A statistical approach for extracting confinement forces and diffusivity from a single blurred trajectory. *Phys Rev E* 93, 053303 (2016).

34. Terasaki, M., Song, J., Wong, J. R., Weiss, M. J. & Chen, L. B. Localization of endoplasmic reticulum in living and glutaraldehyde-fixed cells with fluorescent dyes. *Cell* 38, 101–108 (1984).

35. Lee, C. & Chen, L. B. Dynamic behavior of endoplasmic reticulum in living cells. *Cell* 54, 37–46 (1988).

36. Terasaki, M., Chen, L. B. & Fujiwara, K. Microtubules and the endoplasmic reticulum are highly interdependent structures. *J Cell Biology* 103, 1557–1568 (1986).

37. Lee, C. & Chen, L. B. Endoplasmic Reticulum. *Subcell Biochem* 21, 343–352 (1993).

38. Terasaki, M. & Reese, T. S. Interactions among endoplasmic reticulum, microtubules, and retrograde movements of the cell surface. *Cell Motil Cytoskel* 29, 291–300 (1994).

39. Waterman-Storer, C. M., Gregory, J., Parsons, S. F. & Salmon, E. D. Membrane/microtubule tip attachment complexes (TACs) allow the assembly dynamics of plus ends to push and pull membranes into tubulovesicular networks in interphase Xenopus egg extracts. *J Cell Biology* 130, 1161–1169 (1995).

40. Waterman-Storer, C. M. & Salmon, E. D. Endoplasmic reticulum membrane tubules are distributed by microtubules in living cells using three distinct mechanisms. *Curr Biol* 8, 798–807 (1998).

41. Nehls, S. *et al.* Dynamics and retention of misfolded proteins in native ER membranes. *Nat Cell Biol* 2, 288–295 (2000).

42. York, A. G. *et al.* Instant super-resolution imaging in live cells and embryos via analog image processing. *Nat Methods* 10, 1122–1126 (2013).

43. Li, D. *et al.* ADVANCED IMAGING. Extended-resolution structured illumination imaging of endocytic and cytoskeletal dynamics. *Sci New York N Y* 349, aab3500 (2015).

44. Guo, Y. *et al.* Visualizing Intracellular Organelle and Cytoskeletal Interactions at Nanoscale Resolution on Millisecond Timescales. *Cell* 175, 1430-1442.e17 (2018).

45. Chen, S., Novick, P. & Ferro-Novick, S. ER network formation requires a balance of the dynamin-like GTPase Sey1p and the Lunapark family member Lnp1p. *Nat Cell Biol* 14, 707–716 (2012).

46. Hu, J. *et al.* A Class of Dynamin-like GTPases Involved in the Generation of the Tubular ER Network. *Cell* 138, 549–561 (2009).

47. Renvoisé, B. *et al.* Reep1 null mice reveal a converging role for hereditary spastic paraplegia proteins in lipid droplet regulation. *Hum Mol Genet* 25, 5111–5125 (2016).

48. Chang, J., Lee, S. & Blackstone, C. Spastic paraplegia proteins spastizin and spatacsin mediate autophagic lysosome reformation. *J Clin Investigation* 124, 5249–62 (2014).

49. Liu, T. Y. *et al.* Cis and trans interactions between atlastin molecules during membrane fusion. *Proc National Acad Sci* 112, E1851–E1860 (2015).

50. Wang, S., Tukachinsky, H., Romano, F. B. & Rapoport, T. A. Cooperation of the ER-shaping proteins atlastin, lunapark, and reticulons to generate a tubular membrane network. *Elife* 5, e18605 (2016).

51. Shen, H. *et al.* Single Particle Tracking: From Theory to Biophysical Applications. *Chem Rev* 117, 7331–7376 (2017).

52. Diezmann, A. von, Shechtman, Y. & Moerner, W. E. Three-Dimensional Localization of Single Molecules for Super-Resolution Imaging and Single-Particle Tracking. *Chem Rev* 117, 7244–7275 (2017).

53. Manzo, C. & Garcia-Parajo, M. F. A review of progress in single particle tracking: from methods to biophysical insights. *Rep Prog Phys* 78, 124601 (2015).

54. Chenouard, N. *et al.* Objective comparison of particle tracking methods. *Nat Methods* 11, 281–289 (2014).

55. Beheiry, M. E. *et al.* A Primer on the Bayesian Approach to High-Density Single-Molecule Trajectories Analysis. *Biophys J* 110, 1209–15 (2016).

56. Masson, J.-B. *et al.* Mapping the energy and diffusion landscapes of membrane proteins at the cell surface using high-density single-molecule imaging and Bayesian inference: application to the multiscale dynamics of glycine receptors in the neuronal membrane. *Biophys J* 106, 74–83 (2014).

57. Parutto, P., Heck, J., Heine, M. & Holcman, D. Biophysics of high density nanometer regions extracted from super-resolution single particle trajectories: application to voltage-gated calcium channels and phospholipids. *Sci Rep-uk* 9, 18818 (2019).

58. Kim, S., Leal, S. S., Halevy, D. B., Gomes, C. M. & Lev, S. Structural Requirements for VAP-B Oligomerization and Their Implication in Amyotrophic Lateral Sclerosis-associated VAP-B(P56S) Neurotoxicity*. *J Biol Chem* 285, 13839–13849 (2010).

59. Hua, R. *et al.* VAPs and ACBD5 tether peroxisomes to the ER for peroxisome maintenance and lipid homeostasis. *J Cell Biol* 216, 367–377 (2017).

60. Prosser, D. C., Tran, D., Gougeon, P.-Y., Verly, C. & Ngsee, J. K. FFAT rescues VAPA-mediated inhibition of ER-to-Golgi transport and VAPB-mediated ER aggregation. *J Cell Sci* 121, 3052–3061 (2008).

61. Teuling, E. *et al.* Motor Neuron Disease-Associated Mutant Vesicle-Associated Membrane Protein-Associated Protein (VAP) B Recruits Wild-Type VAPs into Endoplasmic Reticulum-Derived Tubular Aggregates. *J Neurosci* 27, 9801–9815 (2007).

62. Suzuki, H. *et al.* ALS-linked P56S-VAPB, an aggregated loss-of-function mutant of VAPB, predisposes motor neurons to ER stress-related death by inducing aggregation of co-expressed wild-type VAPB: Detailed characterization of VAPB/ALS8. *J Neurochem* 108, n/a-n/a (2010).

63. Yamanaka, T., Nishiyama, R., Shimogori, T. & Nukina, N. Proteomics-Based Approach Identifies Altered ER Domain Properties by ALS-Linked VAPB Mutation. *Sci Rep-uk* 10, 7610 (2020).

64. Papiani, G. *et al.* Restructured endoplasmic reticulum generated by mutant amyotrophic lateral sclerosis-linked VAPB is cleared by the proteasome. *J Cell Sci* 125, 3601–3611 (2012).

65. Calderon, C. P., Weiss, L. E. & Moerner, W. E. Robust hypothesis tests for detecting statistical evidence of two-dimensional and three-dimensional interactions in single-molecule measurements. *Phys Rev E* 89, 052705 (2014).

66. Persson, F., Lindén, M., Unoson, C. & Elf, J. Extracting intracellular diffusive states and transition rates from single-molecule tracking data. *Nat Methods* 10, 265–269 (2013).

67. Monnier, N. *et al.* Inferring transient particle transport dynamics in live cells. *Nat Methods* 12, 838–840 (2015).

68. Schwantes, C. R., McGibbon, R. T. & Pande, V. S. Perspective: Markov models for long-timescale biomolecular dynamics. *J Chem Phys* 141, 090901 (2014).

69. Grimm, J. B. *et al.* A general method to improve fluorophores for live-cell and single-molecule microscopy. *Nat Methods* 12, 244–50, 3 p following 250 (2015).

70. Grimm, J. B. *et al.* Bright photoactivatable fluorophores for single-molecule imaging. *Nat Methods* 13, 985–988 (2016).

71. Grimm, J. B. *et al.* A general method to fine-tune fluorophores for live-cell and in vivo imaging. *Nat Methods* 14, 987–994 (2017).

72. Calderon, C. Data-Driven Techniques for Detecting Dynamical State Changes in Noisily Measured 3D Single-Molecule Trajectories. *Molecules* 19, 18381–18398 (2014).

73. Calderon, C. P. & Bloom, K. Inferring Latent States and Refining Force Estimates via Hierarchical Dirichlet Process Modeling in Single Particle Tracking Experiments. *Plos One* 10, e0137633 (2015).
